# Extinction risk, reconstructed catches, and management of chondrichthyan fishes in the Western Central Atlantic Ocean

**DOI:** 10.1101/2022.01.26.477854

**Authors:** Brendan S. Talwar, Brooke Anderson, Cristopher G. Avalos-Castillo, María del Pilar Blanco-Parra, Alejandra Briones, Diego Cardeñosa, John K. Carlson, Patricia Charvet, Charles F. Cotton, Zoe Crysler, Danielle H. Derrick, Michael R. Heithaus, Katelyn B. Herman, Olga Koubrak, David W. Kulka, Peter M. Kyne, Oscar M. Lasso-Alcalá, Paola A. Mejía-Falla, Jorge Manuel Morales-Saldaña, Beatriz Naranjo-Elizondo, Andrés F. Navia, Nathan Pacoureau, Juan C. Peréz-Jiménez, Riley A. Pollom, Cassandra L. Rigby, Eric V.C. Schneider, Nikola Simpson, Nicholas K. Dulvy

## Abstract

Chondrichthyan fishes are among the most threatened vertebrates on the planet because many species have slow life histories that are outpaced by intense fishing. The Western Central Atlantic Ocean, which includes the greater Caribbean, is a hotspot of chondrichthyan biodiversity and abundance, but is historically characterized by extensive shark and ray fisheries and a lack of sufficient data for effective management and conservation. To inform future research and management decisions, we analyzed patterns in chondrichthyan extinction risk, reconstructed catches, and regulations in this region. We summarized the extinction risk of 180 sharks, rays, and chimaeras using contemporary IUCN Red List assessments and found that over one-third (35.6%) were assessed as Vulnerable, Endangered, or Critically Endangered largely due to fishing. Reconstructed catches from 1950 to 2016 reached their peak in 1992, then declined by 40.2% through the end of the series. The United States, Venezuela, and Mexico were responsible for most catches and hosted large proportions of the regional distributions of threatened species; these countries therefore held the greatest responsibility for chondrichthyan management. The abundance and resolution of fisheries landings data were poor in much of the region, and national-level regulations varied widely across jurisdictions. Deepwater fisheries represent an emerging threat, although many deepwater chondrichthyans currently find refuge beyond the depths of most fisheries. Regional collaboration as well as effective and enforceable management informed by more complete fisheries data, particularly from small-scale fisheries, are required to protect and recover threatened species and ensure sustainable fisheries.

## 1. INTRODUCTION

Fishing has outpaced the slow life histories of many sharks and their relatives (class Chondrichthyes, hereafter ‘sharks and rays’; Cortés, 2000; Worm et al., 2013) and has led to an estimated one-third (37.5%) of sharks and rays being threatened with extinction (Dulvy et al., 2021a). Oceanic sharks and rays present a striking example; between 1970 and 2018, an 18-fold increase in relative fishing pressure reduced their global abundance by 71% (Pacoureau et al., 2021). Sharks inhabiting coral reefs are similarly threatened, with fishing likely responsible for sharks being absent from almost 20% of reefs surveyed globally (MacNeil et al., 2020). The depletion of shark and ray populations could lead to ecosystem-level consequences (Burkholder et al., 2013; Estes et al., 2016; Ferretti et al., 2010) because many of these fishes are apex or mesopredators that range widely and may affect ecosystem processes through predation and associated risk effects, competition, nutrient transport, and bioturbation (Flowers et al., 2021; Heithaus et al., 2008, 2010; Heupel et al., 2014).

Increased concern for fisheries impacts on sharks and rays in recent decades gave rise to numerous initiatives developed to stem or reverse population declines at the national and international level (Shiffman & Hammerschlag, 2016). In 1991, for example, the International Union for Conservation of Nature (IUCN) Species Survival Commission (SSC) Shark Specialist Group (SSG) was founded to promote the sustainable use and conservation of sharks and rays (Fowler et al., 2005), and, in 1993, the United States implemented its Fishery Management Plan for sharks in the Atlantic Ocean (NMFS, 1993). Additionally, in the late 1990s, the United Nations (UN) Food and Agriculture Organization (FAO) developed the International Plan of Action for Conservation and Management of Sharks (IPOA–Sharks), which recommended countries create and implement their own National Plans of Action for sharks and rays (NPOA– Sharks; FAO, 1999). Other management measures (e.g., trade restrictions) were introduced over the next twenty years, but their full implementation is a challenge (Lawson & Fordham, 2018), and their effectiveness remains to be demonstrated on a global scale (Davidson et al., 2016).

In the wider Caribbean, robust shark and ray management is lacking (Davidson et al., 2016; Fowler et al., 2005), and any existing management has been described as a patchwork of inconsistent measures (Kyne et al., 2012). Further, the wider Caribbean was recently one of the most data-deficient regions for sharks and rays in the world (Dulvy et al., 2014). According to the IUCN Red List of Threatened Species (hereafter ‘IUCN Red List’) in 2012, nearly half (47%) of the region’s shark and ray species were assessed as Data Deficient and nearly one in five (19%) were assessed in a threatened category, primarily due to overfishing (Kyne et al., 2012). Some historical accounts and archaeological data suggest that fishing had depleted large marine vertebrates in the Caribbean even before modern fishing technology and scientific research expanded in the mid-1900s (Jackson et al., 2001; McClenachan et al., 2006; Wing & Wing, 2001), although these conclusions are debated (e.g., see Baisre, 2010; McClenachan et al., 2010). As recently as the 1950s, however, sharks were still described as highly abundant (Viele, 1996; Ward-Paige et al., 2010), possibly illustrating the ‘shifting baselines’ concept (Pauly, 1995).

Contemporary trends in shark abundance in the wider Caribbean have been derived from time-series catch data from fisheries-independent surveys and United States-based fisheries (including the pelagic longline fleet that covers much of the Caribbean). These data suggest declines in the abundance or size of some coastal (Cortés et al., 2002; Hayes et al., 2009; McClenachan, 2009) and oceanic sharks (Baum & Blanchard, 2010; Cortés et al., 2007; Jiao et al., 2009), particularly following intense fishing in the 1980s (Bonfil, 1997; Castro, 2013; Musick et al., 1993). The magnitudes of some widely-reported declines in the region’s shark abundance are debated (see Baum et al., 2003; Baum and Myers, 2004; Burgess et al., 2005). Fisher surveys (Graham, 2007) and spatial variation in relative abundance also suggest fishing caused declines in some coastal shark populations – abundance is often highest in heavily managed exclusive economic zones (EEZs; MacNeil et al., 2020), marine reserves (Bond et al., 2012; MacNeil et al., 2020), shark sanctuaries (Clementi et al., 2021), and remote areas far from human population centers (Ward-Paige et al., 2010). There are, however, signs of recent stability and/or recovery in some better-studied shark populations in the United States (Carlson et al., 2012; Peterson et al., 2017), The Bahamas (Hansell et al., 2018; Talwar et al., 2020), and Belize (Bond et al., 2017), largely due to targeted management that began in the 1990s (Castro, 2013; Ward-Paige, 2017). Otherwise, a lack of data has challenged the assessment of shark population trends.

Ray (superorder Batoidea) population trends are poorly known in the wider Caribbean and, for coastal species, trends vary spatially. Precipitous declines in sawfish (Pristidae) abundance are well documented across the entire region, for example (Bonfil et al., 2017; Fernandez-Carvalho et al., 2014; Thorson, 1982), but at least one highly managed, well-studied population of Smalltooth Sawfish (*Pristis pectinata*, Pristidae) is stable and likely recovering in the United States (Brame et al., 2019). Diver observations from 1994 to 2007 suggest that Yellow Stingray (*Urobatis jamaicensis*, Urotrygonidae) abundance declined on coral reefs but increased in some areas where predator populations were overfished (e.g., Jamaica; Ward-Paige et al., 2011). Important ray (and shark) habitats such as coral reef, seagrass, and mangrove ecosystems (White & Sommerville, 2010) have also been degraded in the wider Caribbean (Jackson et al., 2014; Polidoro et al., 2010; Waycott et al., 2009), which can lead to range contractions and increased extinction risk (Yan et al., 2021).

Chimaera (i.e., ghost shark, order Chimaeriformes) population trends are unknown in the wider Caribbean, but chimaeras typically reside offshore, are caught as bycatch, and have little commercial value (Finucci et al., 2021). Globally, their contribution to total chondrichthyan catch is very low (Dulvy et al., 2014). Further, chimaeras primarily reside at depths beyond the maximum depth of most Caribbean fisheries (Finucci et al., 2021). Their populations, along with the populations of deepwater sharks and rays, are probably stable as a result (Dulvy et al., 2014), but remain understudied.

Recently, there have been efforts to reduce data deficiency and improve management for sharks and rays in the region. In 2017, the FAO Western Central Atlantic Fishery Commission (WECAFC), a regional fisheries advisory body that hosts members that fish or are located in FAO Major Fishing Area 31 (Western Central Atlantic, ‘WCA’) and the northern part of FAO Major Fishing Area 41 (Southwest Atlantic), convened the first meeting of the working group on shark and ray conservation and management. The working group highlighted the need to coordinate national and regional management and made several specific recommendations regarding shark and ray fisheries (WECAFC, 2018). It also reviewed a Regional Plan of Action (RPOA–Sharks), a regionally tailored version of the IPOA–Sharks meant to facilitate collaboration in research, data collection, and management. Formal adoption of the RPOA– Sharks was intended for early 2020 (WECAFC, 2019), but it remains in draft form at the time of this writing.

To inform future research and upcoming management decisions, we summarize updated global assessments of shark and ray extinction risk for species found in the WCA using data from the IUCN SSC SSG’s Global Shark Trends Project (Kyne et al., 2020; Dulvy et al., 2021a). We analyze extinction risk according to taxonomy, maximum depth of occurrence, and trophic position. We then examine key threats, particularly fishing, and review current shark and ray management at the international and country (states and territories) level.

## 2. MATERIALS AND METHODS

### 2.1 Application of the IUCN Red List Categories and Criteria

Twenty regional experts and members of the IUCN SSC SSG met for five days at the Cape Eleuthera Institute in Eleuthera, The Bahamas in June 2019. The IUCN Red List Categories and Criteria (Version 3.1) were applied to 113 species of sharks and rays following the Guidelines for Using the IUCN Red List Categories and Criteria (IUCN, 2012; IUCN Standards and Petitions Subcommittee, 2019). Assessments were conducted at the global level (i.e., for the entire global population of each species). Data were collated on the taxonomy, distribution, population status, habitat and ecology, major threats, use and trade, and conservation measures for each species from peer-reviewed literature, fisheries statistics, grey literature, and consultation with species and fisheries experts. For details on each of the eight IUCN Red List Categories and the five Criteria used to assess each category of extinction risk, see Mace et al. (2008), IUCN (2012), and IUCN Standards and Petitions Subcommittee (2019). Briefly, a species is Extinct (EX) when no individuals remain alive and Extinct in the Wild (EW) when it only survives in captivity or in naturalized populations outside its previous range. Critically Endangered (CR) species face an *extremely high risk* of extinction in the wild, Endangered (EN) species face a *very high risk* of extinction in the wild, and Vulnerable (VU) species face a *high risk* of extinction in the wild. These CR, EN, and VU species are considered threatened. Near Threatened (NT) species are close to qualifying or are likely to qualify for a threatened category in the future, and Least Concern (LC) species are widespread or abundant taxa not currently qualifying for, nor close to qualifying for, a threatened category. Data Deficient (DD) species lack sufficient information on either their distribution or population status to adequately assess their extinction risk, and could potentially be LC, CR, or any Category in-between.

Draft assessments were prepared in the IUCN Species Information Service online database, then reviews were solicited from at least two experts trained in applying the IUCN Red List Categories and Criteria with knowledge of the species and fisheries at hand. A summary of the assessments was also provided to the entire IUCN SSC SSG (174 members) for their consult and input prior to submission to the IUCN Red List Unit (Cambridge, UK) for further review and quality checks. Assessments were then published on the IUCN Red List (version 2021-1, www.iucnredlist.org; IUCN, 2021; see Data S3, Dulvy et al., 2021a). The assessments drafted at this workshop made up the majority of those included in this study; the remainder were conducted in the same manner at workshops elsewhere (e.g., oceanic species were assessed during a 2018 workshop in Dallas, Texas in the southern United States).

### 2.2 Geographic & taxonomic scope

The WCA extends from the eastern coast of French Guiana (5°00’N latitude) to the southeastern coast of the United States (36°00’N latitude). It includes the Atlantic Ocean, Gulf of Mexico, and Caribbean Sea from the east coast of North, Central, and South America to 40°00’W longitude (Figure 1; FAO, 2021). It includes waters attributed to 13 continental states, 13 island states, and over 20 territories (associated with Colombia, France, the Netherlands, United Kingdom, and United States), encompassing 14.6 million km^2^.

**Figure 1:**
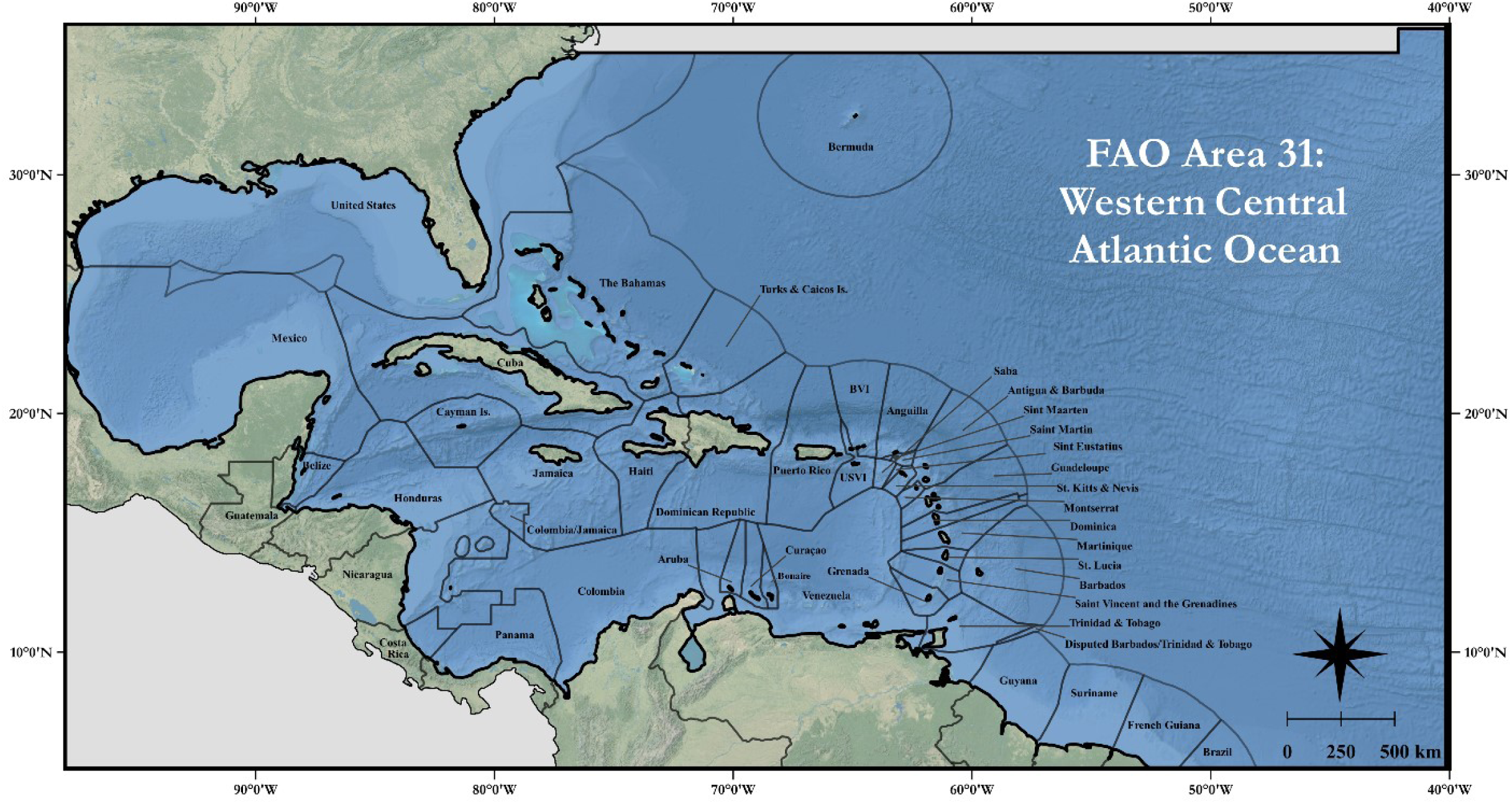
Map of United Nations Food and Agriculture Organization (FAO) Major Fishing Area 31 in the Western Central Atlantic Ocean. National boundaries (Flanders Marine Institute, 2019) are in dark grey and other FAO Areas are shaded grey. Map base layer source: Esri®

We included all marine chondrichthyans assessed on the IUCN Red List that occur in the WCA, including residents and migrants. We excluded freshwater chondrichthyans because their fisheries and management are separate from marine fishes and focused our narrative less on chimaeras and oceanic sharks than other groups because they were evaluated in recent publications (Finucci et al., 2021; Pacoureau et al., 2021). We used the nomenclature and authorities listed in the online Catalog of Fishes (Eschmeyer et al., 2017) and revisions of *Sharks of the World* (Ebert et al., 2013, 2021) for sharks and chimaeras and *Rays of the World* (Last et al., 2016) for rays. We used only global assessments, all of which were available online (www.iucnredlist.org; IUCN, 2021). This review therefore reports the global status of species occurring in the WCA rather than a region-specific assessment, although we note that the assessments of endemic species are limited to the WCA.

### 2.3 Analyzing habitat, trophic level, and threat data

We coded each species according to the IUCN Major Threats and Habitats Classification Schemes (http://www.iucnredlist.org/technical-documents/classification-schemes/habitats-classification-scheme-ver3 and http://www.iucnredlist.org/technical-documents/classification-schemes/threats-classification-scheme) (Salafsky et al., 2008). Species were assigned to one or more of the following habitat classifications: deep benthic, oceanic, neritic, wetlands, intertidal, and coastal/supratidal according to their known depth distribution. We extracted the maximum depth of each species’ depth distribution from the IUCN Red List assessments and compared it across categories of extinction risk. We also extracted trophic level estimates from FishBase (Froese & Pauly, 2021) for each species, then compared trophic levels across categories of extinction risk. We attempted to analyze these data with linear models, but model residuals failed the Shapiro-Wilk test of normality even after data transformation, so we used a non-parametric Kruskal-Wallis test and a post-hoc Dunn’s test to detect differences in both cases. We accounted for multiple comparisons by adjusting p-values using the Benjamini-Hochberg method. Lastly, we coded threats to each species as either present or absent and summarized those threats for all species and then for threatened species only.

### 2.4 Species distributions and conservation responsibility

We mapped the distribution of chondrichthyans in the WCA using IUCN Red List species distribution shapefiles that were built according to taxonomic records summarized in FAO species catalogues (see Dulvy et al., 2014; Dulvy et al., 2021a), *Rays of the World* (Last et al., 2016), revisions of *Sharks of the World* (Ebert et al., 2013, 2021), and recent capture data, expert input, and species checklists (e.g., Mejía-Falla et al., 2019; Tavares, 2019; Weigmann, 2016). Ranges were clipped to the minimum and maximum depth of each species. We set the maximum depth for species without a known depth range to the maximum confirmed depth of the family. We produced a species richness map for all sharks and rays by counting the number of polygons where species distribution maps overlapped. We then used natural neighbor interpolation to interpolate between counts and clipped the output to exclude land. Due to imperfections in the underlying data, these counts should be interpreted for broadscale patterns only. Maps were created with QGIS3 (www.qgis.org).

We estimated jurisdiction-specific conservation responsibility (CoR) to highlight the jurisdictions with the greatest responsibility for conserving globally threatened sharks and rays within the WCA as follows: we assigned threat scores to each species according to their extinction risk, where LC was assigned a zero, NT a one, VU a two, EN a three, and CR a four. No species were assessed as EX or EW. For each jurisdiction (including all countries as well as international waters), we multiplied the threat score of every species present by its proportional range within the WCA in that jurisdiction (Kyne et al., 2020; Rodrigues et al., 2014). We took the sum of those values for each jurisdiction to calculate raw CoR values, then normalized them from 0 to 1 to compare CoR across jurisdictions (where a 1 was assigned to the country with the highest CoR). We then produced a map displaying CoR using Jenks natural breaks classification, which reduces within-class variance and maximizes between-class variance.

### 2.5 Reconstructed fisheries catch data

We extracted reconstructed catch data from the Sea Around Us Project database (www.seaaroundus.org) to examine trends in shark and ray catches from 1950 to 2016 (Pauly et al., 2020). The Sea Around Us database provides estimates of unreported catches (e.g., discards, subsistence, recreational, and small-scale catches) combined with official figures reported by member countries to the UN FAO (Zeller et al., 2016). We used data for the functional groups ‘small to medium sharks ≤ 90 cm’, ‘large sharks ≥ 90 cm’, ‘small to medium rays ≤ 90 cm’, and ‘large rays ≥ 90 cm’ within FAO Area 31 only and then examined patterns in catches over time by fishing entity (i.e., country) and taxonomy (Pauly & Zeller, 2015). Many countries in the WCA have EEZs that extend beyond FAO Area 31, but we did not include catches from those areas (e.g., southern Brazil or the Pacific coast of Central American countries). We did include catches from foreign fleets (e.g., Spain) that occurred in the area.

### 2.6 Management

We collated the most recent stock assessment results for sharks and rays in the WCA from the International Commission for the Conservation of Atlantic Tunas (ICCAT; https://www.iccat.int/Documents/Meetings/Docs/2017_SCRS_REP_ENG.pdf) and the United States’ National Oceanic and Atmospheric Administration (https://www.fisheries.noaa.gov/national/population-assessments/fishery-stock-status-updates).

Assessments indicate a status of ‘overfishing’, ‘overfished’, or ‘unknown’, where overfishing refers to fishing mortality or total catch compromising a stock’s capacity to continuously produce maximum sustainable yield, overfished refers to a stock having a low population size that threatens its ability to reach maximum sustainable yield, and unknown refers to a stock that lacks definitions of overfishing and/or overfished or lacks the data to make a determination.

We estimated jurisdiction-specific Chondrichthyan Management Responsibility (CMR) to reconcile CoR with historical shark and ray fishing and current shark and ray management. The holistic CMR can identify countries that are responsible for high catches of threatened species while rewarding for management in an attempt to highlight 1) countries that may have a high CoR but very low historical catches of sharks and rays and therefore perceive no need for robust management, and 2) countries that may have no modern fisheries for sharks and rays because previous fishing already depleted local populations, leading to limited management where it is urgently required. We calculated CMR using the equation:

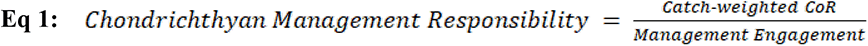

where 1) catch-weighted CoR is a country’s raw CoR (non-normalized) multiplied by its total reconstructed catch (metric tons; mt) of sharks and rays from 1950 to 2016, and 2) Management Engagement (ME) is a country’s percent engagement (0 to 100%) with thirteen management tools (assigned present or absent). These tools were the following:

- Fishing and Finning (3 tools): a ban on shark fishing; a ban on ray fishing; a ban on finning (e.g., a requirement to land fins with associated carcasses or naturally attached)
- UN FAO Plans (2 tools): NPOA–Sharks or RPOA–Sharks, UN FAO National or Regional Plan of Action to Prevent, Deter and Eliminate Illegal, Unreported and Unregulated (IUU) Fishing (NPOA–IUU or RPOA–IUU)
- Other Regulations (1 tool): a single category that included time/area closures, a ban on exports or imports of shark or ray products, species-specific measures, or gear restrictions relevant to sharks and rays
- Party / Signatory / Cooperator to (7 tools): WECAFC; ICCAT; Convention on International Trade in Endangered Species of Wild Flora and Fauna (CITES); Convention on the Conservation of Migratory Species of Wild Animals (CMS); CMS Memorandum of Understanding – Sharks (CMS MOU – Sharks); Protocol for Specially Protected Areas and Wildlife (SPAW) to the Convention for the Protection and Development of the Marine Environment of the Wider Caribbean Region; Port State Measures to Prevent, Deter and Eliminate Illegal, Unreported and Unregulated Fishing (PSM).

We collected this information by searching the scientific and grey literature, UN FAO documents, and news sources. We relied largely on summaries in other reports (Baker-Médard & Faber, 2020; WECAFC, 2018; Koubrak et al., 2021; Kyne et al., 2012; Ward-Paige, 2017; Ward-Paige & Worm, 2017). Where a country’s status was unclear or incomplete, we contacted in-country representatives for additional information. In few cases, all parties involved were unsure of the status of a country relative to a management tool, in which case we used our best judgement in assigning status. Thus, this summary represents our best effort at collating these data, but it may contain errors, particularly where complex overlap occurs between island, national, and international jurisdictions (e.g., Kingdom of the Netherlands). We note that these 13 management tools are not equivalent, and, in some cases, their presence could lead to unintended negative consequences (Castellanos-Galindo et al., 2021).

We omitted jurisdictions where the underlying data structure did not align across CMR components (e.g., where reconstructed catch data were unavailable) except in the case of Saint Martin, St. Barthelemy, and Sint Maarten, which we grouped. We used the mean of their ME and the sums of their CoR and reconstructed catches in this calculation. We then normalized CMR from 0 to 1 for ease of interpretation, where the larger the CMR, the more *unmitigated* risk and responsibility. We also used linear regression to analyze the relationships between CMR components, where a p-value < 0.05 was considered significant.

## 3. RESULTS

### 3.1 Species diversity

We identified 180 assessed shark and ray species in the WCA, which represent 15% of the 1,199 species assessed in the Global Shark Trends Project (Dulvy et al., 2021a). This included 102 sharks, 72 rays, and 6 chimaeras from 12 orders, 46 families, and 83 genera (Table S1). We identified 66 endemic species (36.7% of all species) and 14 near-endemic species (where a small portion of the species’ range extended into another FAO Area; 7.8% of all species). Species richness was highest along the continental margins of North and South America and lowest in oceanic waters (Figure 2). The neritic assemblage was dominated by carcharhiniforms and myliobatiforms (60.4%; *n* = 58 of 96), the oceanic assemblage was dominated by squaliforms and carcharhiniforms (61.4%; *n* = 35 of 57), and the deep slope was dominated by rajiforms and squaliforms (58.4%; *n* = 59 of 101).

**Figure 2:**
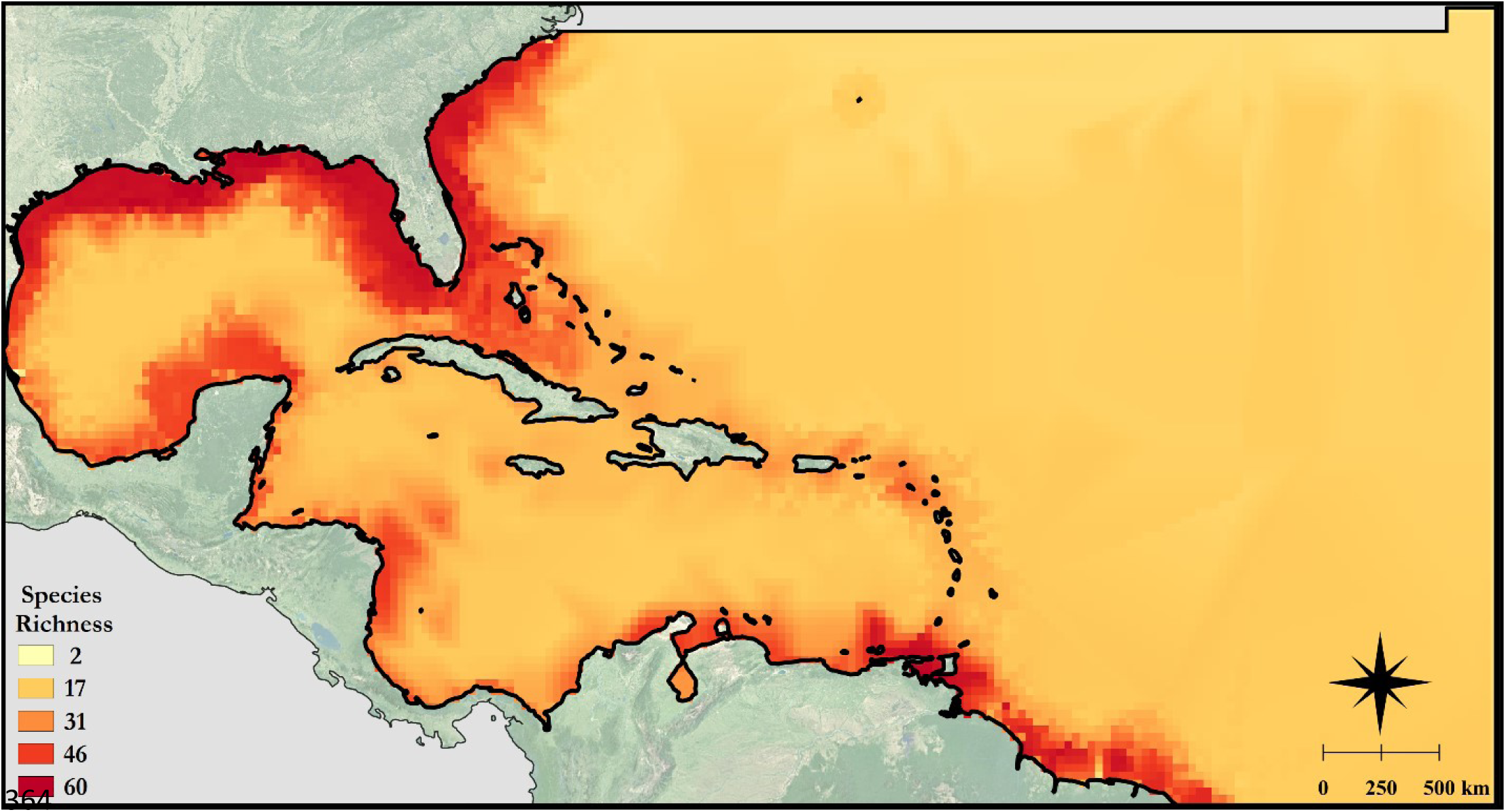
Chondrichthyan species richness in the Western Central Atlantic Ocean based on species distribution maps from the IUCN Red List database (IUCN, 2021). Pixel size is roughly 1025 km^2^. Areas outside of United Nations Food and Agriculture Organization Major Fishing Area 31 are shaded grey. Map base layer source: Esri®

### 3.2 Extinction risk: descriptive patterns in taxonomy, habitat associations, and trophic level

Over one-third (35.6%, *n* = 64 of 180) of all shark and ray species in the WCA were threatened with an elevated risk of extinction (Table 1). Twelve species (6.7%) were Critically Endangered, 25 species (13.9%) were Endangered, and 27 species (15%) were Vulnerable. Seventeen (9.4%) were Near Threatened, over half (53.9%, *n* = 97) of all species were Least Concern, and two (1.1%) species were Data Deficient (Roughskin Spurdog (*Cirrhigaleus asper*, Squalidae) and Carolina Hammerhead (*Sphyrna gilberti*, Sphyrnidae)). All threatened species met Criterion A (‘population reduction measured over the longer of ten years or three generations’) and sub-criterion A2 (‘population reduction observed, estimated, inferred, or suspected in the past where the causes of reduction may not have ceased or may not be understood or may not be reversible’; IUCN, 2012). All NT species nearly met these same criteria. Either sub-criterion A2b (population reduction based on ‘an index of abundance appropriate to the taxon’) or A2d (population reduction based on ‘actual or potential levels of exploitation’; IUCN, 2012) was also cited in each of these assessments. No species met Criterion B (limited geographic range), Criterion C (small population size and decline), Criterion D (very small or restricted population), or Criterion E (quantitative analysis indicating a probability of extinction in the wild exceeding certain thresholds in the future). Out of 180 assessed species, around half (48.9%, *n* = 88) had a decreasing population trend, 8 (4.4%) had an increasing population trend, 70 (38.9%) were listed as stable, and 14 (7.8%) had an unknown population trend.

**Table 1:** The number and percentage of chondrichthyans found in the Western Central Atlantic by IUCN Red List of Threatened Species category. Totals for the threatened categories, which include Critically Endangered, Endangered, and Vulnerable, appear in italics

| **IUCN Red List category** | **Red List status:**<br>**All species (%)** | **Red List status:**<br>**Sharks (%)** | **Red List status:**<br>**Rays (%)** | **Red List status:**<br>**Chimaeras (%)** |
| --- | --- | --- | --- | --- |
| Critically Endangered | 12 (6.7) | 8 (7.8) | 4 (5.6) | 0 (0) |
| Endangered | 25 (13.9) | 15 (14.7) | 10 (13.9) | 0 (0) |
| Vulnerable | 27 (15) | 18 (17.6) | 9 (12.5) | 0 (0) |
| Near Threatened | 17 (9.4) | 11 (10.8) | 5 (6.9) | 1 (16.7) |
| Least Concern | 97 (53.9) | 48 (47.1) | 44 (61.1) | 5 (83.3) |
| Data Deficient | 2 (1.1) | 2 (2) | 0 (0) | 0 (0) |
| *Total threatened* | *64 (35.6)* | *41 (40.2)* | *23 (31.9)* | *0 (0)* |

Contrary to the global trend (Dulvy et al., 2021a), sharks were more threatened than rays in the WCA, with 40.2% (*n* = 41) of sharks and nearly one-third of rays (31.9%, *n* = 23) threatened with extinction (Figure 3). Seven (58.3%) of the twelve orders included at least one threatened species (Figure 4). All species in Rhinopristiformes (100%, *n* = 4) and Orectolobiformes (100%, *n* = 2) were threatened. Over two-thirds of species in Lamniformes (69.2%, *n* = 9) and Myliobatiformes (66.7%, *n* = 16) were threatened. Nearly half (46%, *n* = 23) of the species in Carcharhiniformes, the most speciose order in the WCA, were threatened. Notably, the second most speciose order, Rajiformes, included no threatened species. Of the 45 families in the region, 25 (55.6%) included at least one species in a threatened category.

**Figure 3:**
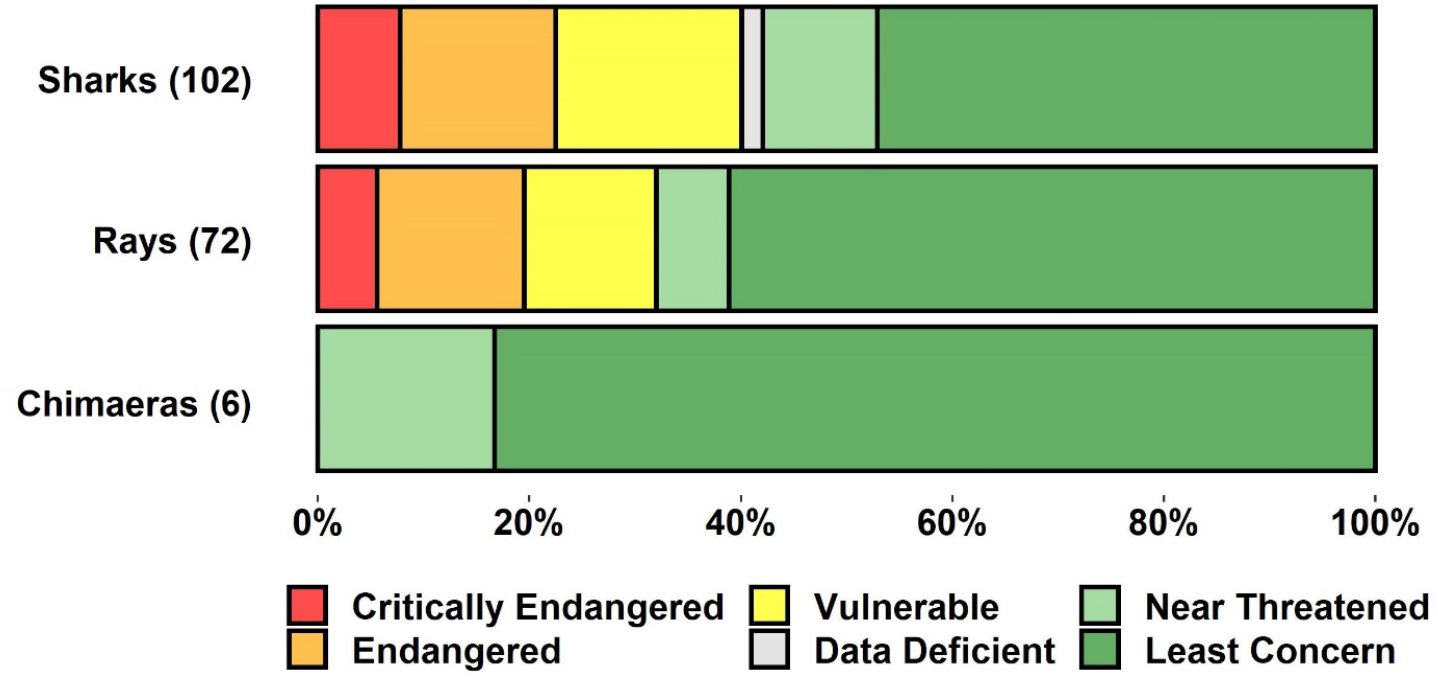
Percentage of sharks, rays, and chimaeras found in the Western Central Atlantic in each IUCN Red List of Threatened Species category. The number of species in each group appears in parentheses

**Figure 4:**
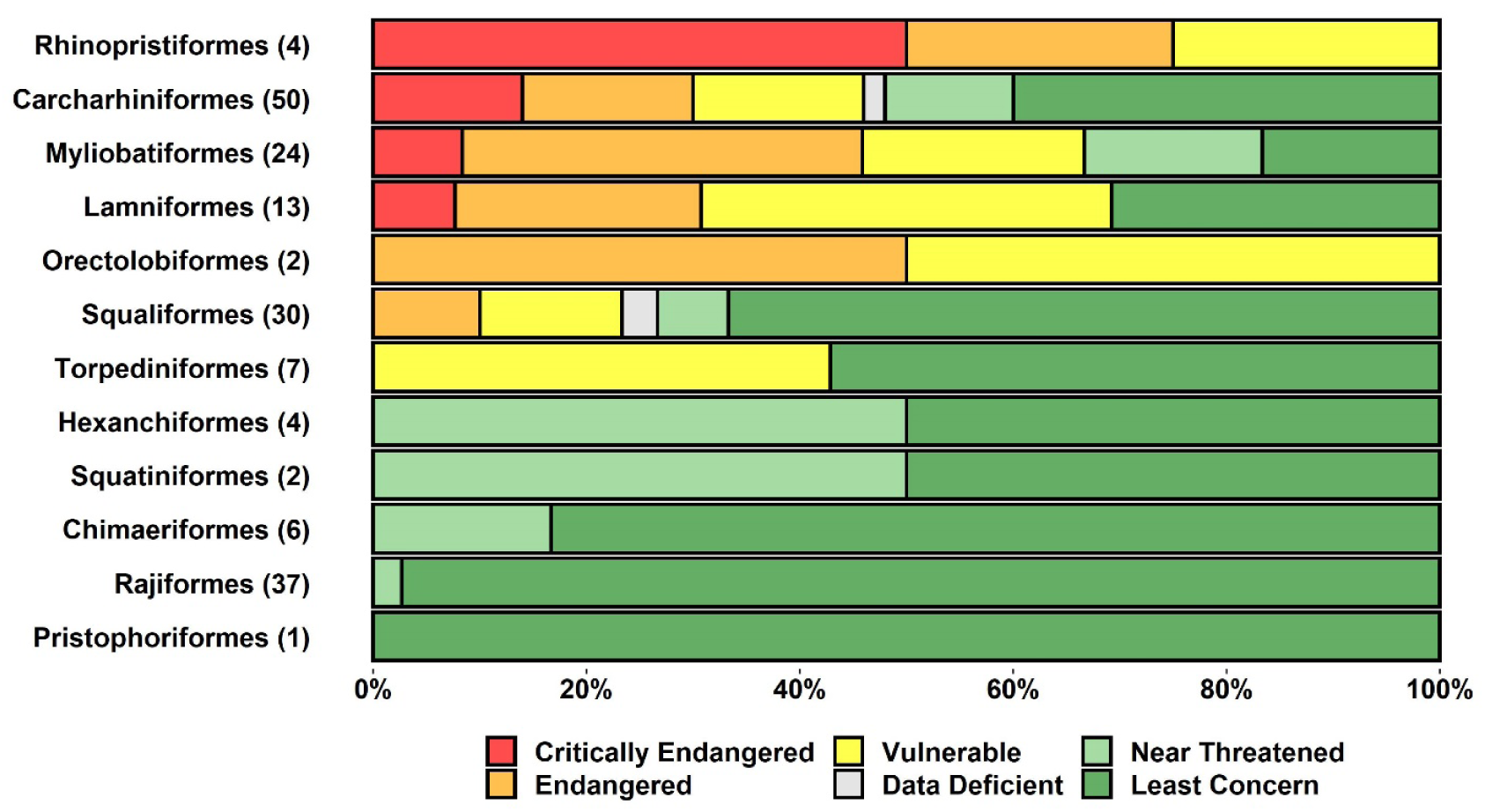
Percentage of each chondrichthyan order found in the Western Central Atlantic by IUCN Red List of Threatened Species category. The number of species in each order appears in parentheses

Sixteen families included only species assessed as LC. Nearly all (95.7%, *n* = 22) species in Rajidae, the most speciose family in the region, were LC. Most (80.4%, *n* = 78) species assessed as LC were associated with depth ranges deeper than 200 m; only 11.9% (*n* = 12 of 101) of species found deeper than 200 m were threatened, and of those the majority (58.3%, *n* = 7 of 12) were assessed as VU. Extinction risk varied with depth (Kruskal-Wallis *x^2^* = 21.06, df = 5, *p* < 0.05), where the maximum depth of LC species (906 ± 588 m; mean ± SD) was significantly greater than the maximum depth of CR (289 ± 390 m; mean ± SD; z = -3.63, *p* < 0.05) and VU species (613 ± 729 m; mean ± SD; z = 2.98, *p* < 0.05; Figure 5). Further, of 78 species with an increasing or stable population trend, 83.3% (*n* = 65 of 78) were associated with the ‘marin deep benthic’ habitat type. There were no differences in trophic levels reported in FishBase among extinction risk categories (Kruskal-Wallis *x^2^* = 6.82, df = 5, *p* = 0.23).

**Figure 5:**
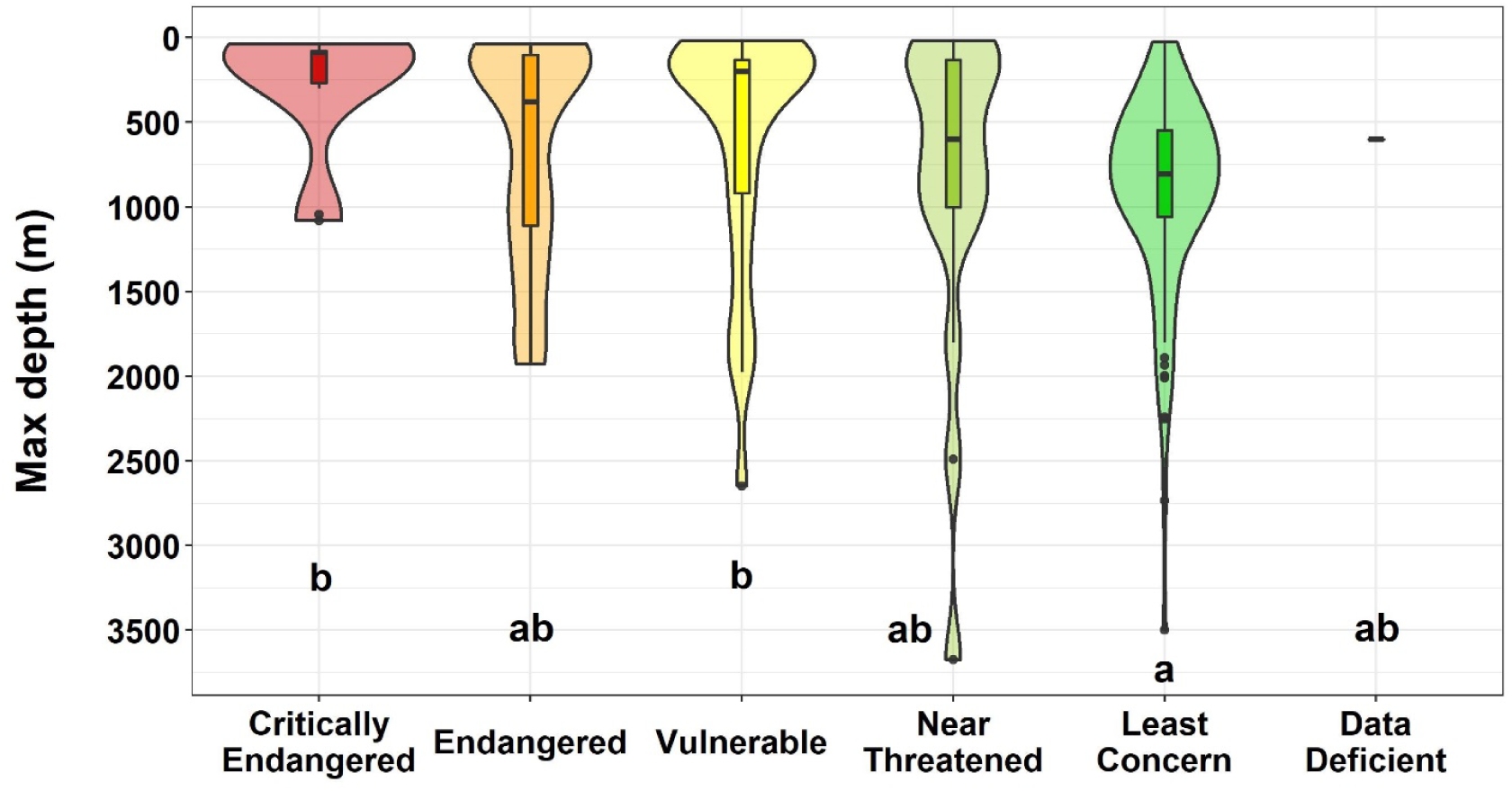
Violin plot of maximum depths of occurrence for all chondrichthyans found in the Western Central Atlantic by IUCN Red List of Threatened Species category. Each dot represents an outlier, horizontal black lines indicate the median, and boxes indicate the interquartile range. Letters represent results of Dunns’s post hoc tests for differences in maximum depth between extinction risk categories, where those sharing the same letter are not significantly different

### 3.3 Endemicity & risk

Of the 66 assessed species endemic to the WCA, 26 were sharks, 36 were rays, and 4 were chimaeras. The top three orders by number of endemic species were Rajiformes (*n* = 29), Carcharhiniformes (*n* = 15), and Squaliformes (*n* = 8). Two-thirds of the chimaeras in the WCA (66.6%; *n* = 4 of 6) were endemic.

No endemic species were assessed as DD. Eighty-nine percent (*n* = 59 of 66) of endemic species were assessed as LC, and 4.5% (*n* = 3 of 66) were assessed as NT. Among sharks, many (72%, *n* = 18 of 25) of the endemic, non-threatened species were lanternsharks (Etmopteridae) and deepwater catsharks (Pentanchidae and Scyliorhinidae). Among rays, many (75.8%, *n* = 25 of 33) were hardnose skates (Rajidae) and pygmy skates (Gurgesiellidae). No endemic chimaeras were in a threatened category, but one endemic shark and three endemic rays were, including the Venezuelan Dwarf Smoothhound (*Mustelus minicanis*, Triakidae; EN), Venezuelan Round Ray (*Urotrygon venezuelae*, Urotrygonidae; EN), Colombian Electric Ray (*Diplobatis colombiensis*, Narcinidae; VU), and Brownband Numbfish (*Diplobatis guamachensis*, Narcinidae; VU). Three near-endemic rays were also threatened – the Painted Dwarf Numbfish (*Diplobatis picta*, Narcinidae; VU), Atlantic Guitarfish (*Pseudobatos lentiginosus*, Rhinobatidae; VU), and Chupare Stingray (*Styracura schmardae*, Potamotrygonidae; EN).

### 3.4 Conservation responsibility

The five countries with the highest conservation responsibility (CoR) were the United States, Venezuela, Mexico, Guyana, and The Bahamas (Figure 6). International waters had the third highest CoR of all jurisdictions (Table S2). Combined, these six jurisdictions accounted for 66.8% of all CoR in the region.

**Figure 6:**
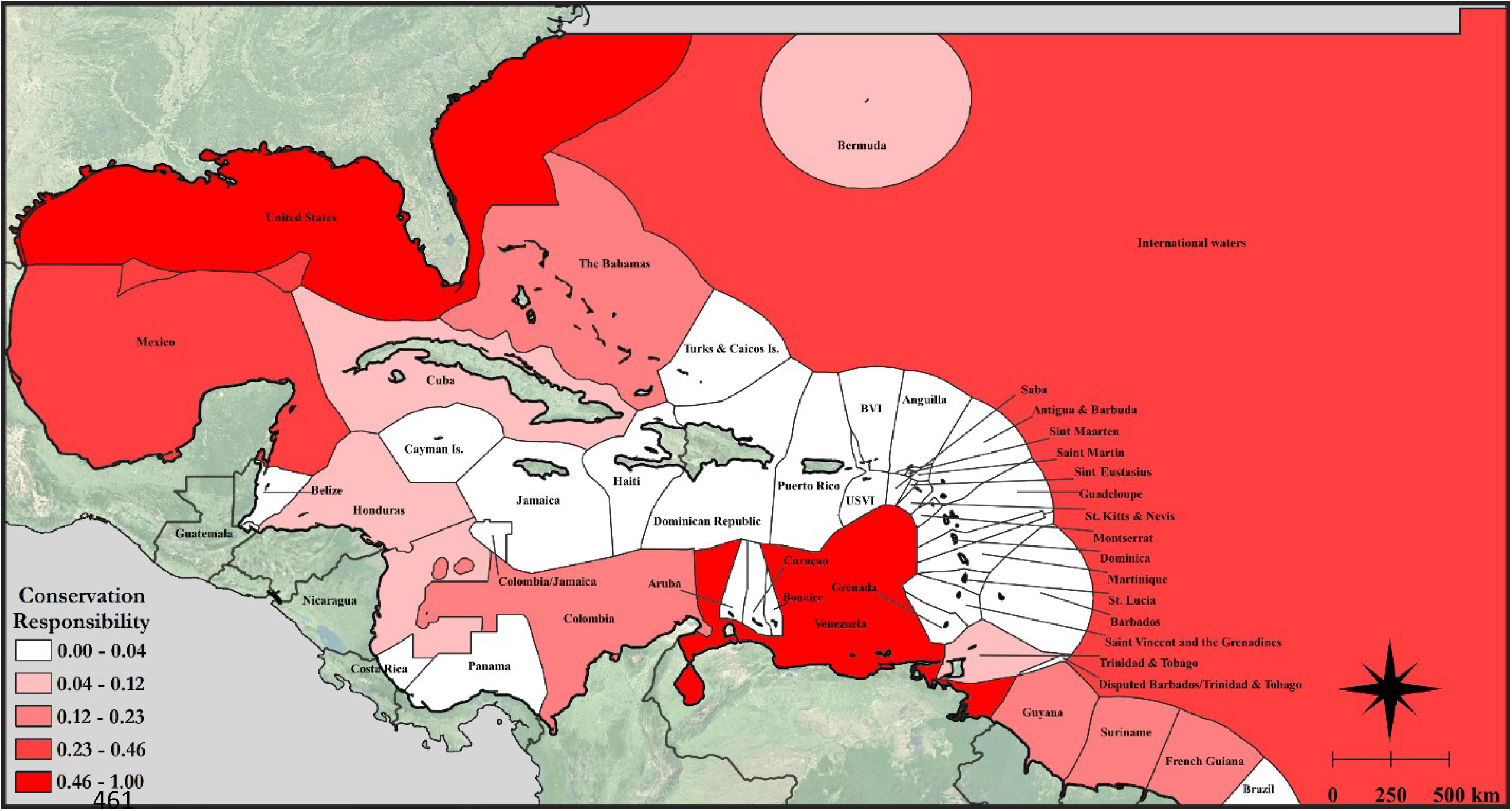
Map of chondrichthyan conservation responsibility for each jurisdiction in the Western Central Atlantic Ocean, where scores are normalized by the maximum score (attributed to the USA) to display from 0 to 1. National boundaries are in dark grey (Flanders Marine Institute, 2019). Regions outside of United Nations Food and Agriculture Organization Major Fishing Area 31 are shaded grey. Map base layer source: Esri®

### 3.5 Key threats

‘Biological resource use’ and, more specifically, ‘fishing and harvesting aquatic resources’, imperiled most sharks and rays (87.8%, *n* = 158 of 180). Threatened species were taken both incidentally and intentionally in large and small-scale fisheries; all threatened species were captured incidentally (100%, *n* = 64 of 64) and most were captured intentionally (81%, *n* = 52 of 64; Figure 7). The threat of overfishing was compounded by habitat loss and degradation and climate change. Habitat loss and degradation imperiled one quarter (26.6%, *n* = 17 of 64) of threatened species primarily through residential and commercial development (and associated habitat modifications), which affected 20.3% (*n* = 13 of 64) of species. Less common pathways to habitat loss and degradation included agriculture and aquaculture (6.3%, *n* = 4 of 64), energy production and mining (4.7%, *n* = 3 of 64), transportation and service corridors (4.7%, *n* = 3 of 64), human intrusions and disturbance (4.7%, *n* = 3 of 64), natural systems modifications (e.g., dams; 1.6%, *n* = 1 of 64), and invasive and other problematic species (1.6%, *n* = 1 of 64). Climate change and severe weather imperiled 14.1% (*n* = 9 of 64) of threatened species. Lastly, pollution (particularly land-based) imperiled 6.3% (*n* = 4 of 64) of threatened species.

**Figure 7:**
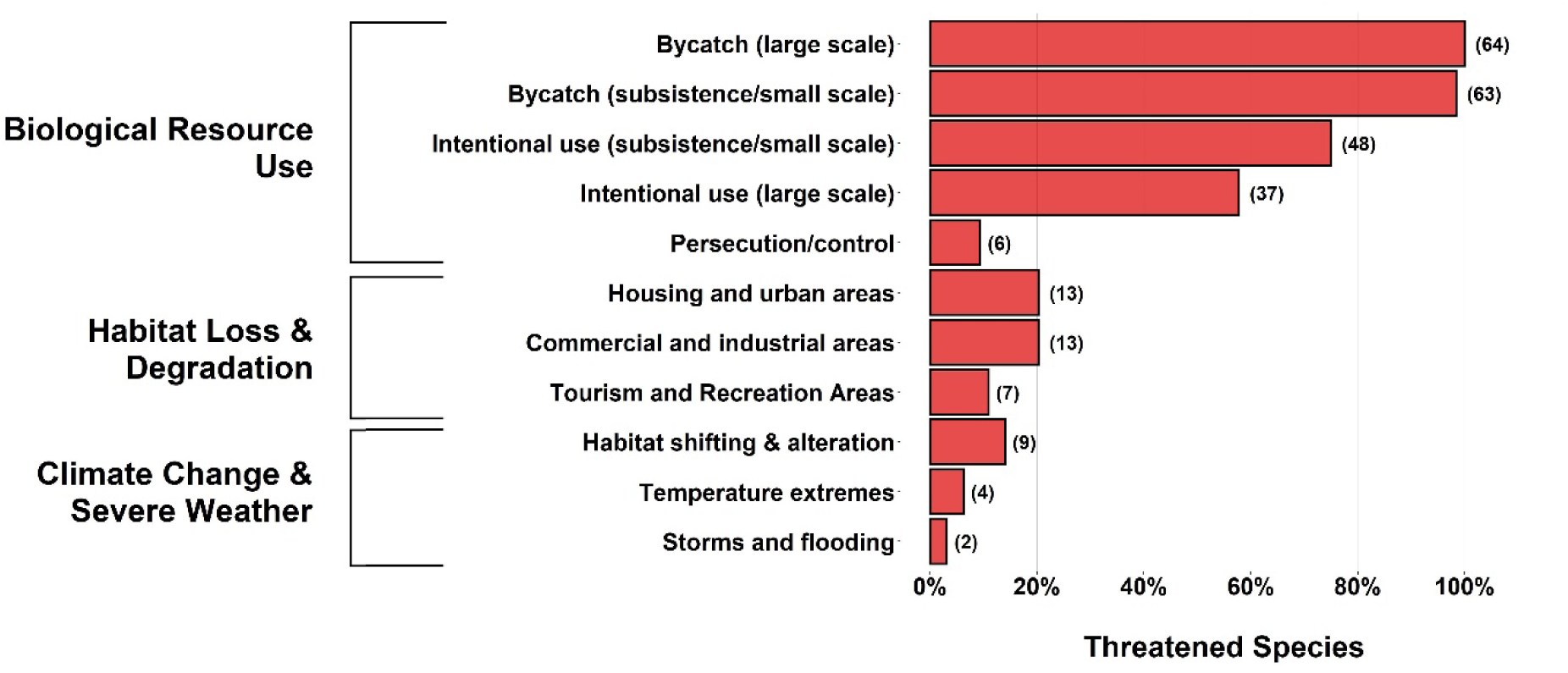
Percentage of threatened sharks and rays in the Western Central Atlantic (n = 64) affected by the most common threats listed in IUCN Red List assessments. The number of species affected by each threat appears in parentheses

### 3.6 Reconstructed fisheries catches

#### 3.6.1 Sharks

Reconstructed shark catches in the WCA more than tripled in 34 years from 1950 (19,458 mt) to 1984 (63,815 mt), plateaued until 1997 (between 48,536 mt and 59,329 mt), then halved over the next decade (2010: 24,015 mt; Figure 8a). In 2011, catches increased to 37,763 mt, due in part to a 451% increase in Venezuelan catches from 2010 to 2011. Spanish catches also rose dramatically from 2009 (0.39 mt) to 2012 (5,701 mt). By 2014, catches of both countries declined to 24.9% of what they were in 2012. By 2016, the total reconstructed catch of sharks in the WCA was half (47.4%) of the peak catch in 1984.

**Figure 8:**
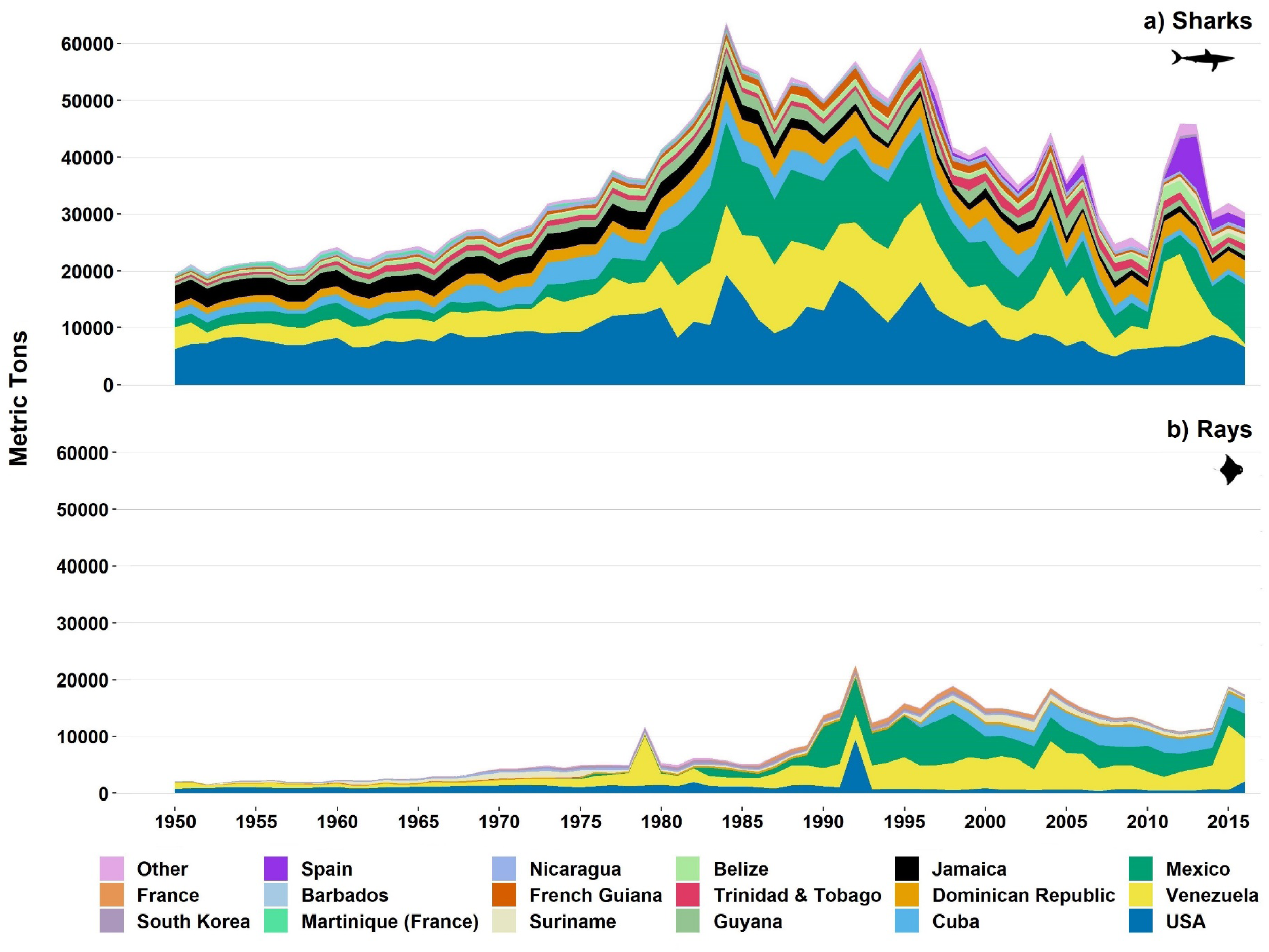
Reconstructed catches of a) sharks and b) rays in the Western Central Atlantic from 1950 to 2016 by country. Those with < 10,000 metric tons of cumulative shark and ray catches across all years are grouped as ‘Other’. Catch data from Pauly et al. (2020) and underlying EEZ boundaries from Claus et al. (2014)

Most shark catches in the region, as well as overall trends in catches, can largely be attributed to fishing by the United States, Venezuela, Mexico, Cuba, the Dominican Republic, and Jamaica (Table 2). Cuba’s maximum annual catch of 4,562 mt occurred in 1977 during a period of elevated catches from 1968 to 2003, when 3,295 mt (± 801 SD) were taken per year. Outside of that period, in 1950 – 1967 and 2004 – 2016, the average annual catch was 1,323 mt (± 343 SD) per year. Jamaica’s maximum annual catch peaked early in 1950 (3,336 mt) and catches declined noticeably from 1978 (3,160 mt) to 1994 (834 mt), then remained low around a mean annual catch of 1,079 mt (± 248 SD). In contrast, catches by the Dominican Republic increased four-fold from a low in 1950 (1,079 mt) to a peak in 1993 (4,390 mt), then remained high around a mean annual catch of 3,277 mt (± 247 SD) through the end of the time series. Foreign fleets were responsible for 2.06% (49,468 mt) of all shark catches.

**Table 2:**
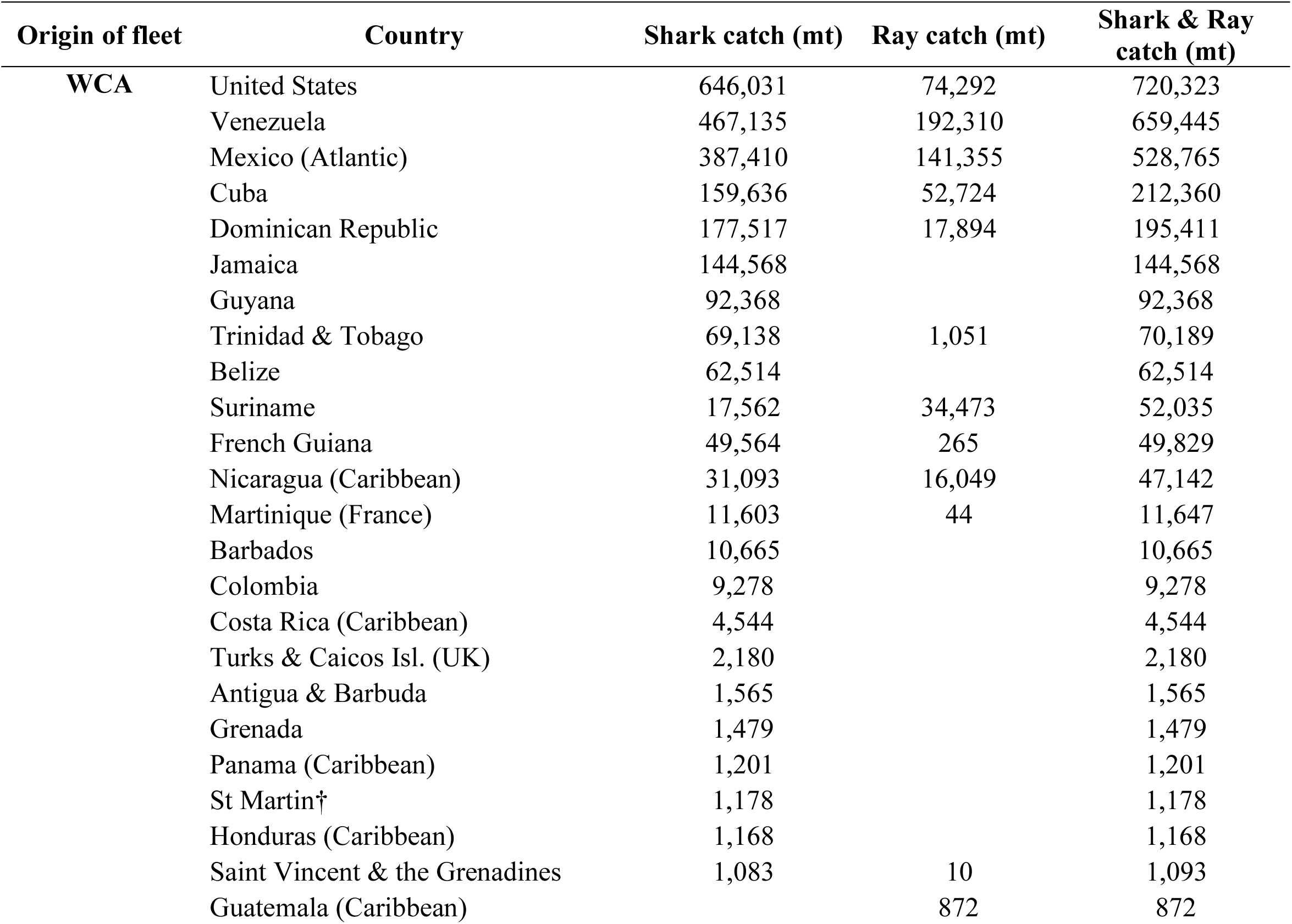
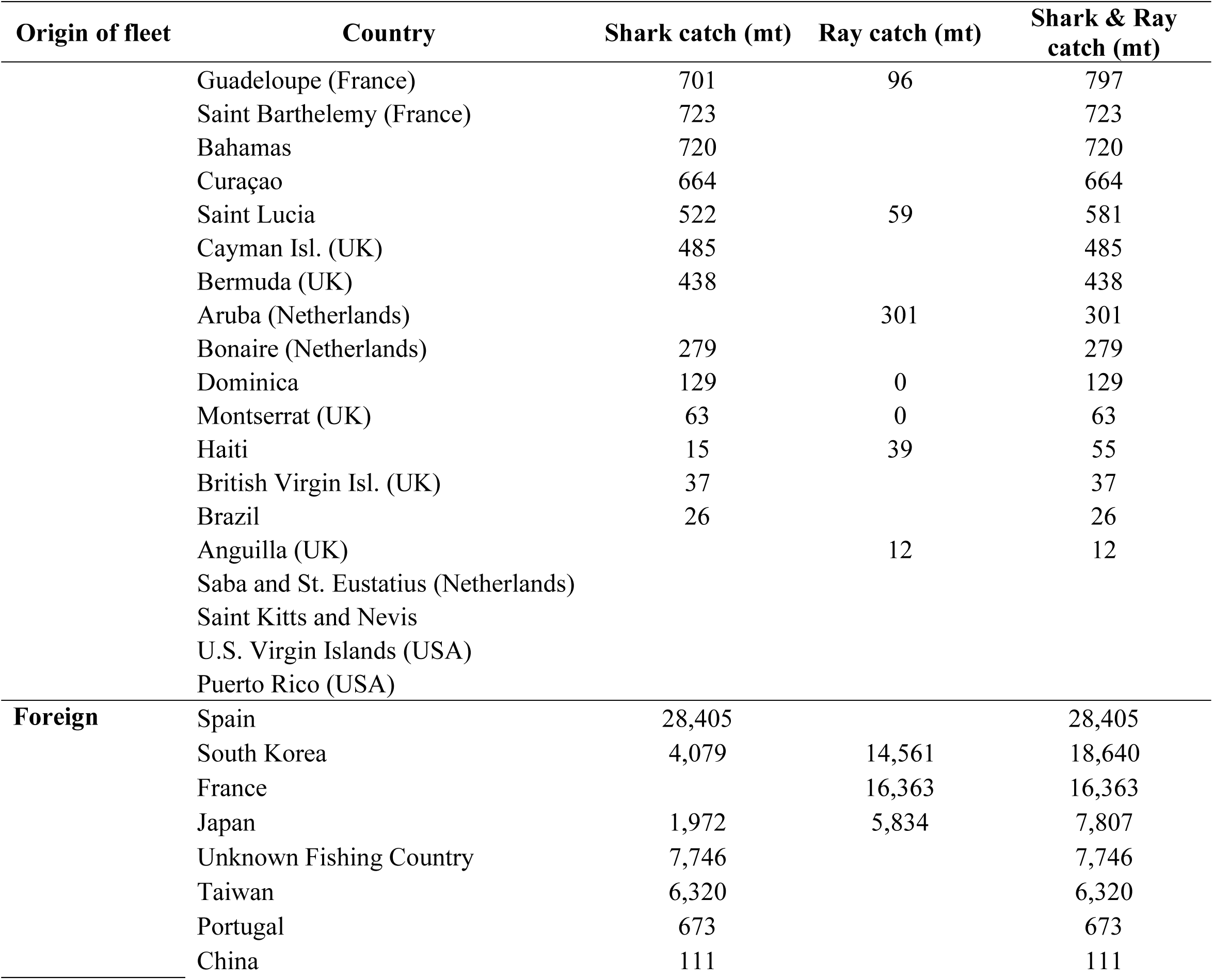
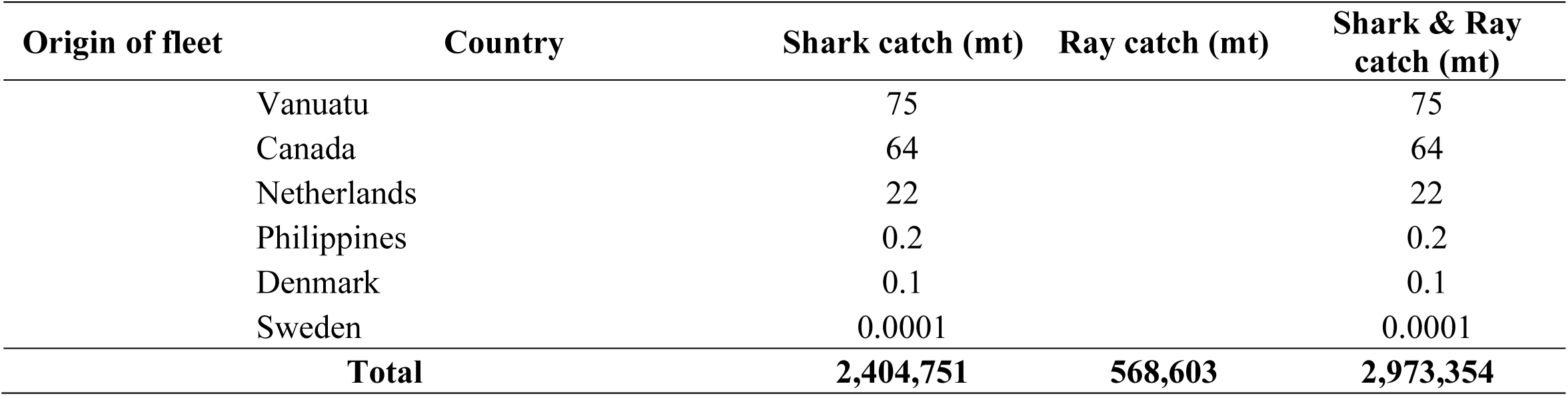
Total reconstructed catch of sharks and rays in the Western Central Atlantic (WCA; FAO Major Fishing Area 31) from 1950 to 2016 by country. Note that each country’s catch outside of the WCA was omitted. Underlying data is from Sea Around Us (Pauly et al., 2020). mt, metric tons

| Origin of fleet | Country | Shark catch (mt) | Ray catch (mt) | Shark & Ray catch (mt) |
| --- | --- | --- | --- | --- |
| WCA | United States | 646,031 | 74,292 | 720,323 |
|  | Venezuela | 467,135 | 192,310 | 659,445 |
|  | Mexico (Atlantic) | 387,410 | 141,355 | 528,765 |
|  | Cuba | 159,636 | 52,724 | 212,360 |
|  | Dominican Republic | 177,517 | 17,894 | 195,411 |
|  | Jamaica | 144,568 |  | 144,568 |
|  | Guyana | 92,368 |  | 92,368 |
|  | Trinidad & Tobago | 69,138 | 1,051 | 70,189 |
|  | Belize | 62,514 |  | 62,514 |
|  | Suriname | 17,562 | 34,473 | 52,035 |
|  | French Guiana | 49,564 | 265 | 49,829 |
|  | Nicaragua (Caribbean) | 31,093 | 16,049 | 47,142 |
|  | Martinique (France) | 11,603 | 44 | 11,647 |
|  | Barbados | 10,665 |  | 10,665 |
|  | Colombia | 9,278 |  | 9,278 |
|  | Costa Rica (Caribbean) | 4,544 |  | 4,544 |
|  | Turks & Caicos Isl. (UK) | 2,180 |  | 2,180 |
|  | Antigua & Barbuda | 1,565 |  | 1,565 |
|  | Grenada | 1,479 |  | 1,479 |
|  | Panama (Caribbean) | 1,201 |  | 1,201 |
|  | St Martin† | 1,178 |  | 1,178 |
|  | Honduras (Caribbean) | 1,168 |  | 1,168 |
|  | Saint Vincent & the Grenadines | 1,083 | 10 | 1,093 |
|  | Guatemala (Caribbean) |  | 872 | 872 |
|  | Guadeloupe (France) | 701 | 96 | 797 |
|  | Saint Barthelemy (France) | 723 |  | 723 |
|  | Bahamas | 720 |  | 720 |
|  | Curaçao | 664 |  | 664 |
|  | Saint Lucia | 522 | 59 | 581 |
|  | Cayman Isl. (UK) | 485 |  | 485 |
|  | Bermuda (UK) | 438 |  | 438 |
|  | Aruba (Netherlands) |  | 301 | 301 |
|  | Bonaire (Netherlands) | 279 |  | 279 |
|  | Dominica | 129 | 0 | 129 |
|  | Montserrat (UK) | 63 | 0 | 63 |
|  | Haiti | 15 | 39 | 55 |
|  | British Virgin Isl. (UK) | 37 |  | 37 |
|  | Brazil | 26 |  | 26 |
|  | Anguilla (UK) |  | 12 | 12 |
|  | Saba and St. Eustatius (Netherlands) |  |  |  |
|  | Saint Kitts and Nevis |  |  |  |
|  | U.S. Virgin Islands (USA) |  |  |  |
|  | Puerto Rico (USA) |  |  |  |
| **Foreign** | Spain | 28,405 |  | 28,405 |
|  | South Korea | 4,079 | 14,561 | 18,640 |
|  | France |  | 16,363 | 16,363 |
|  | Japan | 1,972 | 5,834 | 7,807 |
|  | Unknown Fishing Country | 7,746 |  | 7,746 |
|  | Taiwan | 6,320 |  | 6,320 |
|  | Portugal | 673 |  | 673 |
|  | China | 111 |  | 111 |

| **Origin of fleet** | **Country** | **Shark catch (mt)** | **Ray catch (mt)** | **Shark & Ray catch (mt)** |
| --- | --- | --- | --- | --- |
|  | Vanuatu | 75 |  | 75 |
|  | Canada | 64 |  | 64 |
|  | Netherlands | 22 |  | 22 |
|  | Philippines | 0.2 |  | 0.2 |
|  | Denmark | 0.1 |  | 0.1 |
|  | Sweden | 0.0001 |  | 0.0001 |
|  | **Total** | **2,404,751** | **568,603** | **2,973,354** |

Taxonomic resolution in shark-specific catches was poor; 51.9% of all shark catches were listed only as Elasmobranchii or Chondrichthyes. Much of the regional shark catch from 1950 to 2016 was requiem shark (listed as Carcharhinidae or *Carcharhinus*), which made up 17.7% (426,597 mt) of all catches. Among all recorded shark species, Atlantic Sharpnose Shark (*Rhizoprionodon terraenovae*, Carcharhinidae) made up the largest proportion of catches at 4.5% (109,109 mt), followed by Atlantic Nurse Shark (*Ginglymostoma cirratum*, Ginglymostomatidae; 3.9%, 92,942 mt), Tiger Shark (*Galeocerdo cuvier*, Galeocerdidae; 3.1%, 73,567 mt), Blacktip Shark (*Carcharhinus limbatus*, Carcharhinidae; 2.6%, 62,075 mt), Blue Shark (*Prionace glauca*, Carcharhinidae; 2.1%, 50,505 mt), Bonnethead Shark (*Sphyrna tiburo*, Sphyrnidae; 2.0%, 49,256 mt), and Shortfin Mako (*Isurus oxyrinchus*, Lamnidae; 2.0%, 48,690 mt). From 2012 to 2013, Spain notably caught 14,318 mt of Blue Shark (96% of their total catches of all species during that period). Every other species made up less than 2% of the total catches, although some may be caught in much higher proportions but are difficult to identify at the species level.

#### 3.6.2 Rays

Reconstructed ray catches increased by an order of magnitude from 1950 (2,076 mt) to the peak in 1992 (22,587 mt), then fluctuated between that and a low of 10,892 mt until the end of the series (Figure 8b). Venezuela, Mexico, and the United States were responsible for the largest catches of rays (Table 2). Cuba’s catches increased in the 1990s to contribute substantially to regional catches by 1997 (although national landings data show this increase occurring a decade earlier; PAN-Tiburones, 2015). Catches of rays in the United States were unusually high in 1992 (9,477 mt; 94% of which were stingrays (Dasyatidae)), otherwise they ranged between 408 mt and 2,130 mt. Foreign fleets were responsible for 6.46% (36,758 mt) of all ray catches. As with sharks, taxonomic resolution among recorded ray catches was poor; two-thirds (69%) of all rays were listed as only Batoidea or Rajiformes. The Southern Guitarfish (reported as *Rhinobatos percellens*, now *Pseudobatos percellens*, Rhinobatidae) was caught more than any other listed ray species (73,800 mt, 13% of rays) and is EN.

### 3.7 Management

Some shark and ray species (13.9%, *n* = 25 of 180) were listed on at least one of the following: CITES, CMS, or SPAW. Twenty species were listed on CITES (Appendix I: 2 species, Appendix II: 18 species; Table S1), all of which were also listed on CMS (Appendix I: 11, Appendix II: 9 species). Nine species were listed on SPAW (Annex II: 2, Annex III: 7 species), all of which were also listed on CITES and CMS. Three species were listed on only CMS in Appendix II: Dusky Shark (*Carcharhinus obscurus*, Carcharhinidae; EN), Blue Shark (NT), and Spiny Dogfish (*Squalus acanthias*, Squalidae; VU).

Stock assessments were conducted for the Gulf of Mexico, Atlantic, North Atlantic, or Northwest Atlantic populations of 42 (23.3%, *n* = 42 of 180) shark and ray species that occur in the WCA. Six (14.3%, *n* = 6 of 42) stocks were overfished and eight (19.1%, *n* = 8 of 42) were not overfished (Table S1). Overfishing was occurring in four (9.5%, *n* = 4 of 42) stocks and not occurring in ten (23.8%, *n* = 10 of 42). Twenty-eight (66.7%, *n* = 28 of 42) stocks were assigned an overfished / overfishing status of ‘unknown’.

The type and degree of shark and ray management varied in the WCA (Table 3; see Table S3 for full details and references). The United States had the most detailed management framework that included species-specific catch quotas, time-area closures, gear restrictions, size restrictions, and more. Other countries in the region engaged very little with shark and ray management (e.g., Haiti; Figure 9b). Eleven countries prohibited commercial or all shark (n = 10) or ray (n = 9) fishing, although Honduras’ prohibition on shark fishing included a notable exception for the retention and sale of incidentally caught sharks.

**Figure 9:**
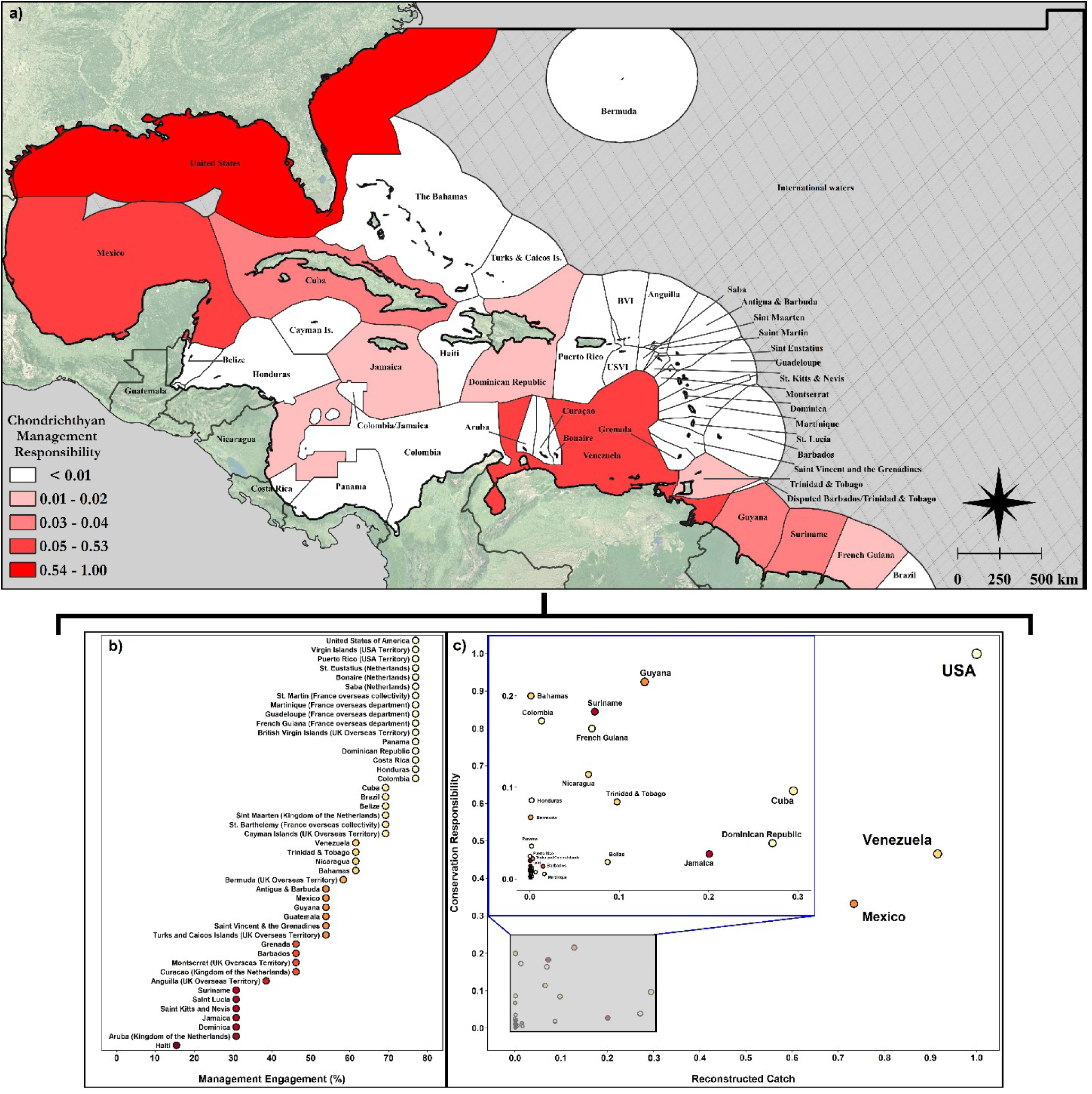
Chondrichthyan Management Responsibility (a) and its components, which include management engagement with thirteen shark and ray management tools (b) and catch-weighted conservation responsibility (c). The Chondrichthyan Management Responsibility Score is normalized by the maximum score (attributed to the USA) to display from 0 to 1. Note that some jurisdictions were omitted and others grouped due to the structure of the underlying data. Catch data from Pauly, Zeller, and Palomares (2020) and underlying EEZ boundaries from Claus et al. (2014)

**Table 3:** Country-level management information, where NPOA–Sharks is a National Plan of Action for the Conservation and Management of Sharks; RPOA–Sharks is a Regional Plan of Action for the Conservation and Management of Sharks; other regulations include time/area closures, a ban on exports of shark or ray products, species-specific measures, or gear restrictions relevant to chondrichthyans; NPOA–IUU is a National plan of action to prevent, deter and eliminate Illegal, Unreported and Unregulated (IUU) Fishing; RPOA–IUU is a Regional plan of action to prevent, deter and eliminate Illegal, Unreported and Unregulated (IUU) Fishing; PSM is the Agreement on Port State Measures; WECAFC is the Western Central Atlantic Fishery Commission; CITES is the Convention on International Trade in Endangered Species of Wild Fauna and Flora; ICCAT is the International Commission for the Conservation of Atlantic Tunas; CMS is the Convention on the Conservation of Migratory Species of Wild Animals; CMS Sharks MoU is the Memorandum of Understanding on the Conservation of Migratory Sharks; and SPAW is the Convention for the Protection and Development of the Marine Environment of the Wider Caribbean Region Specially Protected Areas and Wildlife Protocol. ‘Coop.’ stands for cooperator (a special status in ICCAT with similar rights and obligations to a contracting party) and ‘Sig.’ stands for signatory (where a country has yet to ratify CMS). ‘N/A’ stands for not applicable (where a country is beyond the convention area of an international mechanism). Cells containing a ‘No’ are highlighted in grey. ‘Country’ refers to all states and territories. Full annotated table available in Supporting Information

| Country | Shark<br>fishing<br>ban | Ray<br>fishing<br>ban | Finning<br>ban | NPOA– or<br>RPOA–<br>Sharks¶ | Other<br>regulations | NPOA– or<br>RPOA–IUU | PSM | WECAFC<br>†† | CITES<br>‡‡ | ICCAT<br>§§ | CMS<br>¶¶ | CMS<br>Sharks<br>MoU††† |
| --- | --- | --- | --- | --- | --- | --- | --- | --- | --- | --- | --- | --- |
| Anguilla | No | No | No | No | No | RPOA | No | Yes | Yes | Yes | No | No |
| Antigua & Barbuda | No | No | Yes | NPOA | Yes | NPOA, RPOA | No | Yes | Yes | No | Yes | No |
| Aruba | No | No | No | No | No | RPOA | No | Yes | Yes | No | No | No |
| Bahamas | Yes | No | Yes | No | Yes | RPOA | Yes | Yes | Yes | No | No | No |
| Barbados | No | No | No | No | No | RPOA | Yes | Yes | Yes | Yes | No | No |
| Belize | No | Yes | Yes | NPOA‡, RPOA | Yes | NPOA, RPOA | No | Yes | Yes | Yes | No | No |
| Bermuda | No | No | Yes | No | No | RPOA | No | Yes | Yes | Yes | Yes | Yes |
| Bonaire | Yes | Yes | Yes | No | Yes | RPOA | No | Yes | Yes | No | Yes | Yes |
| Brazil | No | No | Yes | NPOA | Yes | RPOA | No | Yes | Yes | Yes | Yes | Yes |
| British Virgin<br>Islands | Yes | Yes | Yes | No | Yes | RPOA | No | Yes | Yes | Yes | Yes | Yes |
| Cayman Islands | Yes | Yes | Yes | No | No | RPOA | No | Yes | Yes | Yes | Yes | Yes |
| Colombia | Yes | Yes | Yes | NPOA | Yes | RPOA | No | Yes | Yes | No | No | Yes |
| Costa Rica | No | No | Yes | NPOA, RPOA | Yes | RPOA | Yes | Yes | Yes | Coop. | Yes | Yes |
| Cuba | No | No | Yes | NPOA | Yes | RPOA | Yes | Yes | Yes | No | Yes | No |
| Curaçao | No | No | No | No | No | RPOA | No | Yes | Yes | Yes | Yes | No |
| Dominica | No | No | No | No | No | RPOA | Yes | Yes | Yes | No | No | No |
| Dominican<br>Republic | Yes | Yes | Yes | RPOA | Yes | RPOA | No | Yes | Yes | No | Yes | No |
| French Guiana | No | No | Yes | No | Yes | RPOA | Yes | Yes | Yes | Yes | Yes | Yes |
| Grenada | No | No | No | No | No | RPOA | Yes | Yes | Yes | Yes | No | No |
| Guadeloupe | No | No | Yes | No | Yes | RPOA | Yes | Yes | Yes | Yes | Yes | Yes |
| Guatemala | No | No | Yes | NPOA, RPOA | Yes | RPOA | No | Yes | Yes | Yes | No | No |
| Guyana | No | No | Yes | No | No | RPOA | Yes | Yes | Yes | Coop. | No | No |
| Haiti | No | No | No | No | No | RPOA | No | Yes | No | No | No | No |
| Honduras | Yes† | No | Yes | RPOA | Yes | RPOA | No | Yes | Yes | Yes | Yes | No |
| Jamaica | No | No | No | No | No | RPOA | No | Yes | Yes | No | Sig. | No |
| Martinique | No | No | Yes | No | Yes | RPOA | Yes | Yes | Yes | Yes | Yes | Yes |
| Mexico | No | No | Yes | NPOA | Yes | RPOA | No | Yes | Yes | Yes | No | No |
| Montserrat | No | No | No | No | No | RPOA | No | Yes | Yes | Yes | Yes | Yes |
| Nicaragua | No | No | Yes | NPOA, RPOA | No | RPOA | Yes | Yes | Yes | Yes | No | No |
| Panamá | No | No | Yes | NPOA, RPOA | Yes | RPOA | Yes | Yes | Yes | Yes | Yes | No |
| Puerto Rico | No | No | Yes | NPOA | Yes | NPOA, RPOA | Yes | Yes | Yes | Yes | No | Yes |
| Saba | Yes | Yes | Yes | No | Yes | RPOA | No | Yes | Yes | No | Yes | Yes |
| Saint Kitts and<br>Nevis | No | No | No | No | No | NPOA, RPOA | Yes | Yes | Yes | No | No | No |
| Saint Lucia | No | No | No | No | No | RPOA | No | Yes | Yes | No | No | No |

| Country | Shark fishing ban | Ray fishing ban | Finning ban | NPOA– or RPOA– Sharks¶ | Other regulations | NPOA– or RPOA–IUU | PSM | WECAFC †† | CITES ‡‡ | ICCAT §§ | CMS ¶¶ | CMS Sharks MoU††† |
| --- | --- | --- | --- | --- | --- | --- | --- | --- | --- | --- | --- | --- |
| Saint Vincent & the Grenadines | No | No | Yes | No | No | RPOA | Yes | Yes | Yes | Yes | No | No |
| Sint Maarten | Yes | Yes | Yes | No | Yes | RPOA | No | Yes | Yes | No | Yes | No |
| St. Barthelemy | No | No | Yes | No | Yes | RPOA | Yes | Yes | Yes | No | Yes | Yes |
| St. Eustatius | Yes | Yes | Yes | No | Yes | RPOA | No | Yes | Yes | No | Yes | Yes |
| St. Martin | No | No | Yes | No | Yes | RPOA | Yes | Yes | Yes | Yes | Yes | Yes |
| Suriname | No | No | No | No | No | RPOA | No | Yes | Yes | Coop. | No | No |
| Trinidad & Tobago | No | No | Yes | No | No | RPOA | Yes | Yes | Yes | Yes | Yes | No |
| Turks and Caicos Islands | No | No | No | No | Yes | RPOA | No | Yes | Yes | Yes | Yes | Yes |
| United States | No | No | Yes | NPOA | Yes | NPOA, RPOA | Yes | Yes | Yes | Yes | No | Yes |
| Venezuela | No | No | Yes | NPOA | Yes | RPOA | No | Yes | Yes | Yes | No | No |
| Virgin Islands | No | No | Yes | NPOA | Yes | NPOA, RPOA | Yes | Yes | Yes | Yes | No | Yes |
| **Percent Participation** | **22.2** | **20.0** | **71.1** | **35.6** | **60.0** | **100** | **44.4** | **100** | **97.8** | **64.4** | **53.3** | **42.2** |

†Formed in 2011, but modified in 2016 to allow the retention and sale of incidentally caught sharks.
‡NPOA specifically for sharks on the high seas.
§Lawson and Fordham (2018)
¶Note that there is a draft WECAFC Regional Plan of Action that has yet to be adopted. http://www.fao.org/ipoa-sharks/national-and-regional-plans-of-action/en/
††Members: http://www.fao.org/fishery/rfb/wecafc/en#Org-OrgsInvolved
‡‡Parties: https://cites.org/eng/disc/parties/chronolo.php?order=field_official_name&sort=asc
§§Parties: https://iccat.int/en/contracting.html
¶¶Parties: https://www.cms.int/en/parties-range-states
†††Parties: https://www.cms.int/sharks/en/signatories-range-states
‡‡‡Parties: https://www.car-spaw-rac.org/?Who-we-are

Many countries were party to some international agreements, but not others, resulting in a complex matrix of obligations and regulations that in some cases varied even at the island level (e.g., Kingdom of the Netherlands). Of all international management mechanisms, WECAFC had the highest participation (100%), which meant that all countries were also covered by its RPOA–IUU and will be covered by its RPOA–Sharks once it is finalized. Participation in CITES was also high (97.8%); only Haiti was a non-party. The PSMA, a binding agreement that combats IUU fishing, had the lowest participation (44.4%).

Three countries had 92% of Chondrichthyan Management Responsibility (CMR): the United States, Venezuela, and Mexico (Figure 9a). Just ten countries accounted for 99.3% of CMR: the United States, Venezuela, Mexico, Guyana, Suriname, Cuba, Jamaica, French Guiana, Dominican Republic, and Trinidad and Tobago (Table S4). Of those, Suriname and Jamaica had noticeably low Management Engagement (ME) despite having either high Conservation Responsibility (CoR; Suriname) or high historical catches (Jamaica; Figure 9c). There was no relationship between ME and either total reconstructed catch or CoR. However, there was a positive relationship between CoR and total reconstructed catch (*p* < 0.05, adjusted r^2^ = 0.74).

## 4. DISCUSSION

We provide the first comprehensive reassessment of extinction risk for sharks and rays that occur in the WCA and find this region to be a microcosm of the global challenge to their conservation. Thirty-six percent of sharks and rays in the WCA are threatened with an elevated risk of extinction, which is similar to the percentage of sharks and rays threatened globally (Dulvy et al., 2021a). An even larger proportion – nearly half of all sharks and rays in the WCA (48.9%) – exhibit a decreasing population trend across their global range. Overfishing is the overwhelming threat to their populations and has driven declines in all threatened species. The United States, Venezuela, and Mexico overshadow all other countries in the WCA in terms of total reconstructed catch, conservation responsibility, and management responsibility for sharks and rays. National-level regulations and engagement with international management mechanisms vary widely. In light of these findings, we consider patterns in species richness and extinction risk, highlight species of concern, discuss fisheries trends such as finning, the importance of small-scale fisheries, and shrinking refuge at depth, and identify opportunities for improved management.

### 4.1 Species diversity

The WCA is a hotspot of shark and ray biodiversity (Carpenter, 2002; Weigmann, 2016), particularly for endemic (Derrick et al., 2020), evolutionarily distinct (Stein et al., 2018), and deepwater species (e.g., skates; Dulvy et al., 2021a; McEachran & Miyake, 1990). It is comparable to temperate areas of high richness such as the Northeast Atlantic and Southeast Pacific Ocean, but, like coral reef diversity, this Caribbean fauna is only around half as rich as the speciose Indo-West Pacific region (Weigmann, 2016). Species richness in the WCA is highest on the continental shelf, with notably high species richness in large areas of U.S. waters (e.g., along the productive shelf in the Gulf of Mexico) and along the northern coast of South America, particularly at the dynamic boundary between the tropics and subtropics (Dulvy et al., 2014, 2021; Ward-Paige et al., 2010). Longline fishery data suggest high species richness of oceanic sharks along Venezuela’s islands and coast as well as the Guyana shelf, particularly where seasonal upwelling occurs and freshwater from the Orinoco River and Guyanese river drainages meets the Caribbean Sea (Castellanos et al., 2002; Cervigón, 2005; Muller-Karger & Varela, 1990; Tavares & Arocha, 2008). Similarly, marine bony fishes exhibit high species richness along continental Venezuela and Colombia, which could be driven by these same patterns and enhanced by rocky coastlines (Cervigón, 2005; Linardich et al., 2019; Robertson & Cramer, 2014).

We caution that species distributions are best understood in regions with extensive sampling but are still imperfectly known; U.S. waters, for example, exhibit high species richness and simultaneously receive substantial research effort and funding (Linardich et al., 2019; Miloslavich et al., 2010; Robertson & Cramer, 2014). Elsewhere, data gaps are more common, and distributions are particularly challenging to assign to countries in the southern and eastern Caribbean Sea. Deepwater species distributions are data-poor, and records are sometimes limited to a single specimen, which often reflects a lack of deep-sea fisheries and research (e.g., American Pocket Shark (*Mollisquama mississippiensis*, Dalatiidae), Kyne & Herman, 2020a; Campeche Catshark (*Parmaturus campechiensis*, Pentanchidae), Kyne & Herman, 2020b).

### 4.2 Extinction risk

#### 4.2.1 Spatial & temporal comparisons

The proportion of threatened sharks and rays in the WCA is higher today (35.6%) than it was in 2012 (18.5%; Kyne et al., 2012), but is similar to the modern global estimate (32.6 – 45.5%; Dulvy et al., 2021a). This change is largely due to new information being incorporated into species assessments. Only three species had a genuine change (i.e., a real change in the rate of decline, population size, range size, or habitat) in IUCN Red List Category since their last assessment, where the status of all three worsened: Blacknose Shark (*Carcharhinus acronotus*, Carcharhinidae; previously NT, now EN), Night Shark (*Carcharhinus signatus*, Carcharhinidae; previously VU, now EN), and Whale Shark (*Rhincodon typus*, Rhincodontidae; previously VU, now EN). None of these three species are endemic to the WCA, although much of the Blacknose Shark’s range is in this region.

Globally, most threatened sharks and rays occur in coastal shelf waters, particularly in the tropics (Dulvy et al., 2021a); we found the same trend for the subset of WCA species, where CR and VU species occurred significantly shallower than LC species. As such, the bulk of Conservation Responsibility (CoR) fell on the countries with the largest EEZs that included the most coastal, shelf-associated habitats (e.g., the United States, Venezuela, and Mexico) with two exceptions. International waters and The Bahamas had high CoR despite consisting of only oceanic habitats or being an insular nation, respectively. International waters, in particular, cover a large proportion of the distributions of wide-ranging and highly threatened species in the WCA. The Bahamas also includes large expanses of threatened shark and ray habitat, supports high species richness that characterizes the Florida Straits region, and has well-studied sharks and rays.

The WCA was previously one of the most data-deficient regions in the world for sharks and rays (Dulvy et al., 2014). The proportion of DD species dropped from 47% (*n* = 71 of 151 assessed species) in 2012 (Kyne et al., 2012) to just 1.1% (*n* = 2 of 180) in 2021, marking substantial progress in reducing data-deficient blind spots that can lead to flawed species-specific management (Walls & Dulvy, 2020). Seventy-seven species that we included in our review, some of which were not previously recognized in the WCA, were assessed as DD in 2012. Of those, the vast majority (76.6%, *n* = 59 of 77) are now LC and some (7.8%, *n* = 6 of 77) are NT. Eleven (14.3%, *n* = 11 of 77) species formerly assessed as DD are now threatened at the global level, including two CR (Smalltail Shark (*Carcharhinus porosus*, Carcharhinidae) and Scoophead Shark (*Sphyrna media*, Sphyrnidae)), five EN (Bramble Shark (*Echinorhinus brucus*, Echinorhinidae), Lesser Devilray (*Mobula hypostoma*, Mobulidae), Chilean Devilray (*Mobula tarapacana*, Mobulidae), Venezuelan Dwarf Smoothhound, and Chupare Stingray), and four VU species (Bullnose Ray (*Myliobatis freminvillii*, Myliobatidae), Southern Eagle Ray (*Myliobatis goodei*, Myliobatidae), Brazilian Sharpnose Shark (*Rhizoprionodon lalandii*, Carcharhinidae), and Atlantic Nurse Shark). These eleven species need to be recognized and incorporated into management plans in the WCA with an emphasis on the endemic Venezuelan Dwarf Smoothhound and near-endemic Chupare Stingray.

The Roughskin Spurdog is the only previously assessed species that remains DD. It is a poorly known deepwater species (73 – 600 m depth range) that may be caught as bycatch, but the degree to which fishing affects its population is unknown (Finucci et al., 2020). The Carolina Hammerhead is the other modern DD species. It was recently described, is difficult to identify (Quattro et al., 2013), and was assessed as DD because its depth and geographic distribution, and hence interaction with fisheries, could not be determined (VanderWright et al., 2020). Given that all other hammerhead sharks (Sphyrnidae) in the WCA are threatened, however, this status could be masking a high level of extinction risk to the Carolina Hammerhead.

#### 4.2.2 Species of concern

The WCA hosts a number of threatened oceanic sharks (e.g., mackerel sharks (Lamnidae), thresher sharks (Alopiidae), and some requiem sharks (Carcharhinidae)) and rays (e.g., devil rays (Mobulidae)), particularly in the Gulf of Mexico and U.S. Atlantic (Dulvy et al., 2021a; Pacoureau et al., 2021). Fisheries mortality has caused significant population declines in some of these species (e.g., Oceanic Manta Ray (*Mobula birostris*, Mobulidae); Miller and Klimovich, 2017); they are among the most threatened groups of sharks and rays in the region along with hammerheads, sawfishes, guitarfishes (Rhinobatidae), and very large, highly migratory species (e.g., Whale Shark), all of which are recognized as groups of extreme conservation concern (Dulvy et al., 2016; 2021; Pacoureau et al., 2021). These species are prominent on CITES, CMS, and SPAW appendices and annexes, which highlights the need for international cooperation in managing these species and for countries to meet their national-level commitments to these agreements.

Among the four threatened endemics in the WCA, the VU Colombian Electric Ray and VU Brownband Numbfish are considered irreplaceable based on their small ranges (Dulvy et al., 2014). Although they are relatively productive, both species are captured in poorly managed and intense artisanal demersal trawl fisheries throughout their small geographic ranges in Colombia and Venezuela and are suspected to have declined by 30–49% over the past three generations (Pollom, Herman, et al., 2020b; Pollom, Herman, et al., 2020c). The other two endemic threatened species in the WCA are the EN Venezuelan Dwarf Smoothhound and the EN Venezuelan Round Ray. The former is targeted and caught as bycatch in trawl and longline fisheries off Venezuela and Colombia; it was inferred to have declined by > 99% over the past three generations based on declining landings of smoothhounds (Triakidae) in Venezuela (Pollom, Lasso-Alcalá, et al., 2020). The latter is captured in demersal trawl fisheries and artisanal beach seine fisheries in Colombia but is now rarely observed in catches in Venezuela; its population is suspected to have declined by 50–79% in the last ten years (Pollom, Herman, et al., 2020a).

The threatened near-endemic species (Painted Dwarf Numbfish, Atlantic Guitarfish, and Chupare Stingray) are also subject to high fishing pressure in parts of their ranges (Dulvy et al., 2021b; Pollom et al., 2020; Pollom, Charvet, Faria, et al., 2020a). The Painted Dwarf Numbfish is captured in intense demersal trawl fisheries throughout its small range off northern South America from at least as far west as Venezuela to Brazil (Pollom, Charvet, Faria, et al., 2020a) and is considered irreplaceable (Dulvy et al. 2014). It may find some refuge from fishing at depth (Pollom, Charvet, Faria, et al., 2020a). The Atlantic Guitarfish finds some refuge from trawl fisheries in the U.S. Gulf of Mexico but is a common bycatch species in Mexican shrimp trawl fisheries and exposed to intense unmanaged fisheries elsewhere (Pollom et al., 2020). The Chupare Stingray similarly has refuge at the northern part of its range (e.g., The Bahamas) but is subject to high fishing pressure along the coasts of Venezuela, Colombia, the Guianas, and northern Brazil, where it is presently very rare (Dulvy et al., 2021b). Although the Daggernose Shark (*Isogomphodon oxyrhynchus*, Carcharhinidae; CR), and Wingfin Stingray (*Fontitrygon geijskesi*, Dasyatidae; CR) are not near-endemic to the WCA, we consider them ‘irreplaceable’ because of their threatened status and small ranges that extend from eastern Venezuela to the northern coast of Brazil (Dulvy et al., 2014; Pollom, Charvet, Faria, et al., 2020b; 2020c).

Research is required on the life history, distribution, abundance, and fishery interactions of all threatened endemic, near-endemic, and irreplaceable species, the vast majority (77.8%, *n* = 7 of 9) of which are rays. Conservation responsibility for these species falls solely on countries in the WCA, namely Venezuela, Colombia, Suriname, Guyana, French Guiana, and Brazil. We recommend these countries monitor the status and prioritize the management of these species.

### 4.3 Fisheries trends

Shark and ray catches peaked in the WCA (1992) before they peaked globally (2003; Davidson et al., 2016; Pauly et al., 2020), but regional and global trends followed a similar pattern: there was a substantial increase in catches and landings from 1950 to the 1990s/2000s, followed by a period of decline. In the WCA, reconstructed catches declined 40.2% between 1992 and 2016 while overall fishing effort rose in the region by about 1.1% annually after 1950 (Anticamara et al., 2011). Thus, regional catch-per-unit-effort has probably declined by greater than 50% over the equivalent of three generations for many shark and ray species (which would result in a population reduction sufficient for a species to qualify as Endangered), suggesting fishing is driving their extinction risk in the WCA.

#### 4.3.1 Finning

Some of the most intense shark fishing in the WCA occurred from the 1970s to the early 1990s (Bonfil, 1997; Musick et al., 1993) as negative attitudes towards sharks and the demand for and trade in shark fins increased (Castro, 2013; Worm et al., 2013). With increased demand, some local fin prices also rose, even quadrupling in Guatemalan markets by the mid-2000s (Graham, 2007). Numerous countries in the WCA participated in the fin trade (e.g., Guyana, Trinidad and Tobago; Fowler et al., 2005); 21% of CR Scalloped Hammerhead (*Sphyrna lewini*, Sphyrnidae) fins sampled in Hong Kong, for example, came from the western Atlantic (Chapman et al., 2009). But the global volume of fins imported into Hong Kong (i.e., demand) decreased by 2013 (Shea & To, 2017) and was expected to decrease further in both Hong Kong and China in subsequent years (Dent & Clarke, 2015). Fin prices also dropped in some parts of the WCA as the global trade in shark meat products increased 4.5% per year from 2000 to 2011 (Dent & Clarke, 2015). In some places, meat overtook fins as the most profitable shark product (e.g., northeastern Brazil; Martins et al., 2018). By the mid-2010s, the contribution of Scalloped Hammerhead fins from the Southwest Atlantic, Caribbean Sea, and Northwestern Atlantic randomly sampled in Hong Kong markets was roughly 8.5% (Fields et al., 2020). Silky Shark (*Carcharhinus falciformis*, Carcharhinidae) fin trimmings similarly sampled in markets in Hong Kong and mainland China suggested almost no contribution from Atlantic populations (Cardeñosa et al., 2020) despite the Silky Shark being the second most common species in the fin trade at that time (Cardeñosa et al., 2018). These limited insights and a lack of evidence in the literature suggest little contemporary large-scale shark finning (the removal of fins and discarding of its carcass at sea) in the WCA (Kyne et al., 2012), although finning does occur illegally (e.g., finless carcasses are frequently landed at northern Brazilian ports notwithstanding national law; Feitosa et al., 2018). Fins from landed carcasses also enter the fin trade through legal pathways in even the WCA’s most highly managed and developed fisheries (e.g., United States; Dulvy et al., 2017; Ferretti et al., 2020).

#### 4.3.2 The importance of small-scale fisheries & landings data

Even at low levels of effort, small-scale fishing can significantly reduce the biomass of slow-growing fishes such as sharks and rays (Pinnegar & Engelhard, 2008) and affect critical life stages (e.g., juveniles in possible nursery habitats; Tagliafico et al., 2021). In the WCA, the size, economic contribution, and catch of small-scale fleets has been increasing for decades (Baremore et al., 2021; Canty et al., 2019), and overfishing is occurring in nearly double the percentage of small-scale fisheries (46%) as it is in commercial fisheries (28%; Singh-Renton & McIvor, 2015). The significance of small-scale fishing is highlighted by Mexico and Venezuela, which we identified as two of the top three shark and ray fishing nations in the WCA; small-scale fishing boats comprise 97% of the marine fishing fleet in Mexico (Fernández et al., 2011), and artisanal sources supply 94% of the shark catch in Venezuela (Marquez et al., 2019; Tavares, 2019). Yet, the WCA’s small-scale fisheries are managed less intensely than its large-scale commercial fisheries (Singh-Renton & McIvor, 2015), and, for those affecting sharks and rays, small-scale fisheries are poorly known (Kyne et al., 2012) while large-scale fisheries are better-studied (e.g., see SouthEast Data, Assessment, and Review reports, http://sedarweb.org/sedar-projects; Bonfil, 1997; Peterson et al., 2017; Tavares & Arocha, 2008).

The small-scale fisheries impacting sharks and rays in the WCA are heterogeneous and widespread, and their effort and catch are poorly described (Bonfil, 1997). We found surprisingly little information on ray landings in the WCA and stress further monitoring despite few directed ray fisheries in the region (e.g., in the United States, Cuba, and Mexico; Pérez-Jiménez & Mendez-Loeza, 2015; WECAFC, 2018). Further, the WCA’s country-level landings statistics reported to the FAO have very low species-specific resolution (Dulvy et al., 2014; WECAFC, 2018), with over half of shark and ray catches identified as only ‘chondrichthyan’, ‘elasmobranch’, ‘batoid’, or ‘rajiform’. Mexico, despite being the third largest shark and ray fishing country in the WCA, records catches in only three categories – small sharks (< 1.5 m), large sharks (> 1.5 m), and rays (Pérez-Jiménez & Mendez-Loeza, 2015). Venezuela, despite being the second largest shark and ray fishing country in the WCA, recorded sharks and rays as a single category until 1990, then in three groups (miscellaneous sharks, *Mustelus* spp., and miscellaneous rays) until 2007, after which finer level identification was confounded by a lack of training for fisheries monitoring staff (Tavares, 2019). The situation in smaller shark and ray fishing nations is similar; in Guatemala, only two government fisheries staff monitor its entire ∼150 km Caribbean coast, which hinders landings verification (Hacohen-Domené et al., 2020). This poor resolution is not compatible with effective species-specific management. Some recent studies have begun to fill these gaps by monitoring small-scale fisheries landings (e.g., Guyana – Kolmann et al., 2017; Venezuela – Marquez et al., 2019; Panama – Návalo et al., 2021). In the Belizean shark fishery, for example, a new low-cost method of analyzing fisher-contributed secondary shark fins was successful in determining species and size composition of catches and may prove valuable across the WCA in the future (Quinlan et al., 2021). More research on fishing effort, catch, and baseline abundance data is required to assess populations and adapt management priorities (Bizzarro et al., 2009; Kyne et al., 2012; Pérez-Jiménez & Mendez-Loeza, 2015).

#### 4.3.3 Shrinking refuge at depth

Since 1950, global fisheries have increasingly expanded into the deep sea (Morato et al., 2006). In the Atlantic Ocean, deepwater sharks like gulper sharks (Centrophoridae) and kitefin sharks (Dalatiidae) occurring as deep as 1000 m were reported in fisheries landings as early as 1990 (Morato et al., 2006). Although we found many endemic and LC species in the WCA to be associated with deep habitats that can provide refuge from fishing pressure (Dulvy et al., 2014, 2021; Walls & Dulvy, 2021), we note that this refuge may be shrinking as fishing activities continue to develop in the region’s deep waters (Arana et al., 2009; Baremore et al., 2016).

In the WCA, many deepwater habitats (> 200 m) are accessible to small-scale fishers due to the proximity of these habitats to shore, and consequently deepwater sharks and rays are already caught as bycatch and sometimes targeted. Along the MesoAmerican Barrier Reef, for example, this access coupled with declining yields in coastal fisheries led to the emergence of small-scale deepwater fisheries that use longlines, hook and line, traps, and gillnets to target ‘red snappers’ (e.g., Queen Snapper (*Etelis oculatus*, Lutjanidae), Silk Snapper (*Lutjanus vivanus*, Lutjanidae), Blackfin Snapper (*Lutjanus buccanella*, Lutjanidae)) and groupers (e.g., Yellowedge Grouper (*Hyporthodus flavolimbatus*, Serranidae), Misty Grouper (*Hyporthodus mystacinus*, Serranidae)) between 100 and 550 m (Baremore et al., 2021; WECAFC, 2018). Most small-scale deepwater fisheries in the WCA similarly target this snapper and grouper complex. Off Guatemala, fishers catch and discard some small deepwater sharks and chimaeras, while they target or retain others for meat or liver oil (Finucci et al., 2021; Hacohen-Domené et al., 2020; Polanco-Vásquez et al., 2017). In Venezuela, overfishing of shallow water stocks has led to deepwater (200–800 m) fishing north of Isla de Margarita and Paria Peninsula (eastern region, near Trinidad) and along the coast of Falcón (western region, near Aruba), where endemic and near-endemic species of deepwater sharks, rays, and chimaeras are now caught (OM Lasso-Alcalá, *unpublished data*). Deepwater sharks are also targeted in Honduras (Baremore et al., 2016) and caught off Saba Bank (de Graaf et al., 2017), Curaçao (Van Beek et al., 2013), Belize (Quinlan et al., 2021), northern Cuba (Ruiz-Abierno et al., 2021), and the southern Gulf of Mexico (Pérez-Jiménez & Mendez-Loeza, 2015). In the northern Gulf of Mexico, deep reef-fish longline fisheries and shrimp trawl fisheries also catch deepwater sharks as bycatch, most of which are discarded (Scott-Denton et al., 2011; Scott-Denton & Williams, 2013; Zhang et al., 2014), and, in The Bahamas, recreational fishers often catch small deepwater sharks while targeting red snappers with electric reels (BS Talwar, *pers. obs.*). Across these WCA fisheries, the Dusky Smoothhound (*Mustelus canis*, Triakidae; NT), Cuban Dogfish (*Squalus cubensis*, Squalidae; LC), Atlantic Sixgill Shark (*Hexanchus vitulus*, Hexanchidae; LC), Sharpnose Sevengill Shark (*Heptranchias perlo*, Hexanchidae; NT), Night Shark (EN), gulper sharks (*Centrophorus* spp., Centrophoridae; EN where assessed), and some catsharks (Scyliorhinidae; LC) are the most common deepwater species in landings (Baremore et al., 2021; de Graaf et al., 2017; Hacohen-Domené et al., 2020; Marquez et al., 2019; Quinlan et al., 2021; Scott-Denton et al., 2011; Van Beek et al., 2013).

Although many of the WCA’s deepwater sharks and rays are now currently assessed as LC, our knowledge of their biology and ecology remains incredibly limited. These species also typically lack stock assessments (Table S1; Baremore et al., 2021; Kyne & Simpfendorfer, 2010), and many are intrinsically vulnerable to overfishing due to their life histories (García et al., 2008; Simpfendorfer & Kyne, 2009; Rigby & Simpfendorfer, 2015). Thus, a precautionary approach to their management should be emphasized if deepwater fisheries are further developed in the WCA (Simpfendorfer & Kyne, 2009), which some governments appear to be pursuing (e.g., Belize; Baremore et al., 2021; Kyne et al., 2012).

### 4.4 Management opportunities and priorities

The WCA is geopolitically complex, with more maritime boundaries in the Caribbean alone than in any other Large Marine Ecosystem (Martinez et al., 2017). It also contains highly developed, large countries with extensive fisheries management regimes (e.g., United States) alongside economically challenged small island developing states with limited management capacity (e.g., Haiti). Nutrient-rich continental shelves host industrial fisheries while nutrient-poor coral reefs support artisanal fisheries a short distance away (Singh-Renton & McIvor, 2015). It is not surprising that approaches to shark and ray management vary widely in the region and that challenges to improved management and regular stock assessment include consistency and harmonization in data collection, fisheries monitoring, funding, training, and enforcement. Our findings underscore the objectives of the WECAFC RPOA–Sharks in meeting these challenges (WECAFC, 2018).

Chondrichthyan Management Responsibility and its components (catch-weighted CoR and ME) should be interpreted carefully and only within the WCA; a country may have large shark and ray fisheries elsewhere that were not considered in our analysis. Catch-weighted CoR uses reconstructed catch data from the Sea Around Us Project which improves often low-resolution and sometimes incomplete data self-reported by countries to the FAO (Maharaj et al., 2018). Colombia’s reconstructed catch data for sharks and rays is underestimated, for example, because Colombia does not report ray catches from large-scale fisheries, and many years of shark and ray landings data are missing from government records (Caldas et al., 2009). Despite reconstructed catch data for Colombia showing no ray catches from 1950 to 2016, recent data indicate that rays represent 7.2% of the total volume of small-scale fish and invertebrate catches at three locations in the Colombian Caribbean (Squalus Foundation – AUNAP, *unpublished data*). Thus, Colombia’s catch-weighted CoR and non-normalized CMR are underestimated. Still, most fishing in the Colombian Caribbean is small-scale and results in far fewer shark and ray catches than in the WCA’s major shark and ray fishing nations (PA Mejía-Falla, *pers. obs.*). Normalized CMR, while imperfect due to these and similar errors in the underlying data, provides a relative comparison between national-level catches at the best resolution available.

Taken as a relative measure, Chondrichthyan Management Responsibility can provide a blueprint for regional management priorities and leadership. The countries with the highest CMR in the WCA – the United States, Venezuela, and Mexico – have large expanses of nutrient-rich ecosystems along the continental shelf which support their high reconstructed catches and elevate their CoR. Even if these countries were fully engaged with every management mechanism (i.e., had 100% ME), they would still dominate CMR because their catch-weighted CoR is so high relative to other countries. A high CMR does not necessarily indicate current overfishing, however. Despite leading the WCA in CMR (and catch-weighted CoR), the United States currently offers *some* of the best examples of sustainable shark and ray fishing in the world and acts as a refuge for many threatened sharks and rays (Ferretti et al., 2020; Simpfendorfer & Dulvy, 2017), some of which have experienced preliminary recoveries in U.S. waters (Peterson et al., 2017). Alternatively, Mexico and Venezuela host data-poor fisheries where reference points and stock status are largely unknown, and institutional management capacity is lacking (Pérez-Jiménez & Mendez-Loeza, 2015; Tavares, 2019). Mexico, for example, is not a party to SPAW, CMS, or CMS MoU Sharks; it falls in the lower 50% of WCA countries in management engagement despite its very high CMR.

The WCA’s other major historical shark and ray fishing nations (e.g., Cuba, Dominican Republic, Jamaica) and those with high CoR (e.g., Guyana, Suriname, and French Guiana) formed a second set of countries with significant CMR. Jamaica and Suriname stand out as countries requiring improved management given their high CMR but low ME. Haiti’s lack of shark and ray management also requires immediate action. The high CoR of international waters calls attention to the importance of managing highly migratory sharks and rays through international fisheries management bodies (Tavares & Arocha, 2008; Walls & Dulvy, 2021). Given 100% participation of WCA countries in WECAFC, its upcoming RPOA–Sharks provides a unique opportunity to achieve that end, particularly given WECAFC’s broad taxonomic and geographic jurisdiction. In comparison, ICCAT’s jurisdiction is limited to oceanic species caught by fleets targeting tunas and tuna-like fishes (WECAFC, 2018). However, currently WECAFC does not have the authority to adopt binding management measures.

Improved enforcement is required in much of the WCA, particularly in small-scale fisheries (Kyne et al., 2012; Martins et al., 2018; Saavedra-Díaz et al., 2016). Sharks and rays are caught and landed despite protected status in numerous countries (Feitosa et al., 2018; Gallagher et al., 2015; Van Beek et al., 2013). Along Guatemala’s Caribbean coast, limited fisheries patrols and a lack of funding for enforcement have resulted in unregulated fishing in Guatemalan waters and roving bandit dynamics in neighboring EEZs such as Belize and Honduras (Berkes et al., 2006; Graham, 2007; Hacohen-Domené et al., 2020). Shark fins may also move across international borders to be sold in poorly regulated markets (Kyne et al., 2012). Ineffective management and enforcement of marine protected areas (MPAs) is also common (Bustamante et al., 2014; Perera-Valderrama et al., 2018). In addition, extractive activities are allowed in many MPAs; only 0.5% of the protected areas in the Caribbean associated with European Union and UK Overseas Territories prohibit all extractive activities (Martinez et al., 2017). Generally, funding for enforcement is insufficient and the detection of illegal activity is too infrequent to encourage compliance (although it varies by sub-region; Singh-Renton & McIvor, 2015). At the international level, even when a country is party to an international agreement or treaty, it may not have implemented national regulations to meet its commitments (which are sometimes voluntary or non-binding; e.g., IPOA–Sharks, CMS MoU Sharks; Fischer et al., 2012). The following WCA countries, for example, either partially meet or do not meet their mandatory commitments to protected sharks and rays on CMS Appendix I: Antigua and Barbuda, Cuba, Costa Rica, Honduras, Jamaica, Netherlands (Aruba and Curaçao), Panama, and the United Kingdom (Bermuda, Anguilla, Montserrat, and the Turks and Caicos) (Lawson & Fordham, 2018).

Although we focused primarily on fisheries, national priorities can be established using other value frameworks that provide alternative justification for shark and ray management. Shark and ray tourism, for example, can offer a profitable, non-consumptive alternative to fishing for some species and some people (Gallagher & Hammerschlag, 2011; Kyne et al., 2012). The Bahamas provides an example of how a small island developing state without sufficient fisheries management and enforcement (Sherman et al., 2018) is still able to benefit from the non-extractive use of sharks and rays. As a regional leader in shark and ray ecotourism, it boasts the world’s largest shark diving economy, which generates $113.8 million USD annually (Haas et al., 2017). Although The Bahamas has a rich and abundant shark and ray fauna, over 90% of national expenditures from shark dives came from dives focused on the Caribbean Reef Shark (*Carcharhinus perezi*, Carcharhinidae; Haas et al., 2017), which is one of the most abundant and ubiquitous reef-associated sharks in effectively managed areas in the WCA (MacNeil et al., 2020) and also offers tourist appeal in other locations (e.g., Belize; Graham, 2014). The Cayman Islands also offers a long-standing example of successful non-extractive use; ‘Stingray City’, off Grand Cayman, features tens of Southern Stingrays (*Hypanus americanus*, Dasyatidae) that interact with tourists in what may be the oldest example of shark and ray tourism in the world (Ormond et al., 2016). This site plays a major role in ray-specific tourism generating up to $50 million USD annually for the Cayman Islands (Vaudo et al., 2018), while shark-associated diving and non-extractive use generates an additional $46.8 – $62.6 million USD every year (Ormond et al., 2016). Although shark and ray ecotourism is not without its challenges (Gallagher & Huveneers, 2018), under the right circumstances it can have a net conservation and economic benefit (Gallagher et al., 2015) and may be appropriate for countries with low reconstructed catches and high CoR (e.g., Colombia).

### 4.5 Conclusions

Sharks and rays are among the most threatened vertebrates on our planet, second only to the amphibians (Dulvy et al., 2021a). Protecting CR and EN sharks and rays from fishing, particularly endemic and near-endemic species, remains a regional and global priority (Dulvy et al., 2021a). Unmonitored small-scale fisheries in the WCA likely contribute heavily to shark and ray declines and may grow to threaten some shelf-associated deepwater species. Effective and enforceable fisheries management informed by basic species-specific data on abundance and catch is urgently required across the WCA. Managing shark and ray fisheries has the potential to reduce mortality, halt declines, and promote recovery while supporting food security and livelihoods through sustainable fishing of less-threatened species (Booth et al., 2019; Dulvy et al., 2021a). A robust management toolbox is available to achieve that end (Booth et al., 2020; MacNeil et al., 2020), but improved implementation of locally appropriate tools is required (Davidson et al., 2016).

## ACKNOWLEDGEMENTS

We thank all who volunteered their time and expertise to conduct IUCN Red List assessments. We thank E. Brooks, N. Higgs, and the staff of the Cape Eleuthera Institute for hosting the regional workshop. We thank Georgia Aquarium and Al Dove for supporting KBH. NKD was supported by Natural Science and Engineering Research Council, Canada and Canada Research Chairs program. This project was funded by the Shark Conservation Fund, a philanthropic collaborative pooling expertise and resources to meet the threats facing the world’s sharks and rays. The scientific results and conclusions, as well as any views or opinions expressed herein, are those of the author(s) and do not necessarily reflect those of institutions or data providers. This is contribution No. XX from the Institute of Environment at FIU and No. XX from the Exuma Sound Ecosystem Research Project.

## DATA AVAILABILITY STATEMENT

Underlying data are available through the Sea Around Us Project database (www.seaaroundus.org) and IUCN Red List website (www.iucnredlist.org).

---

**Table S1:**
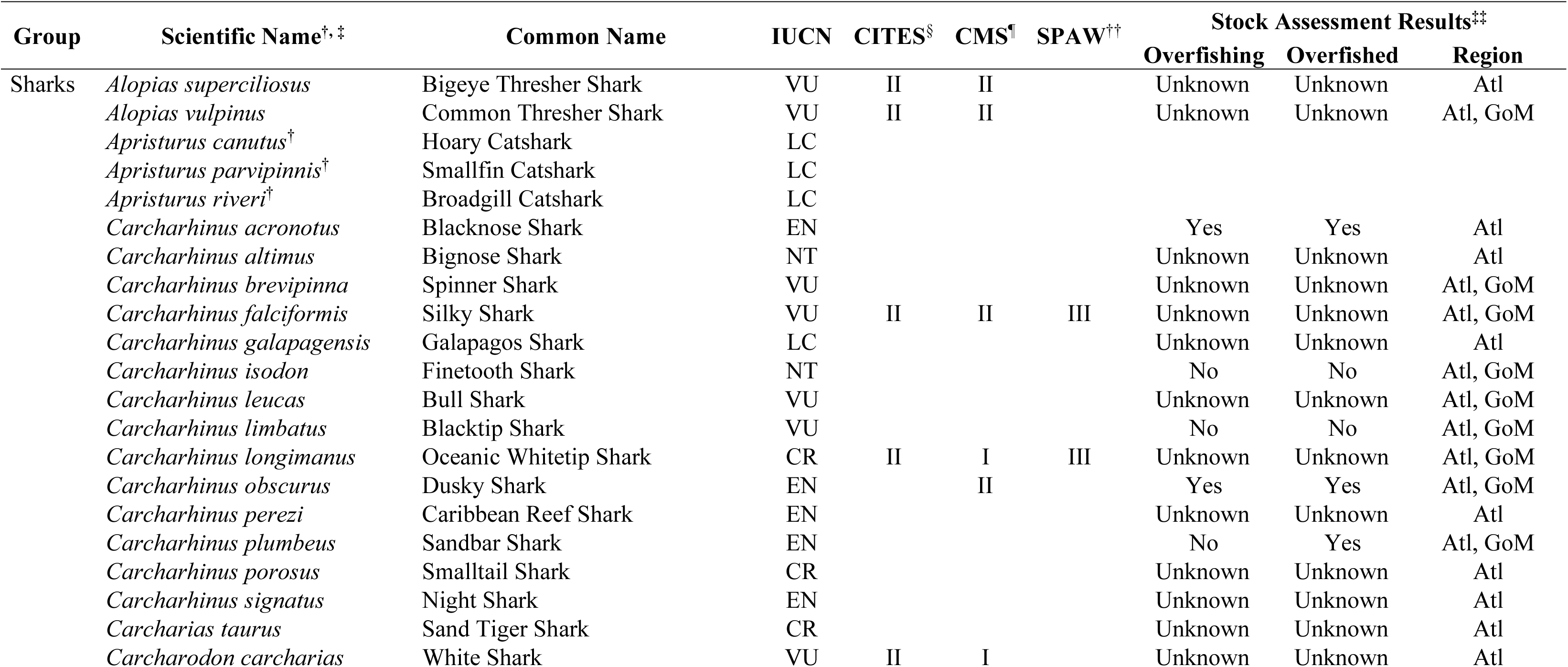
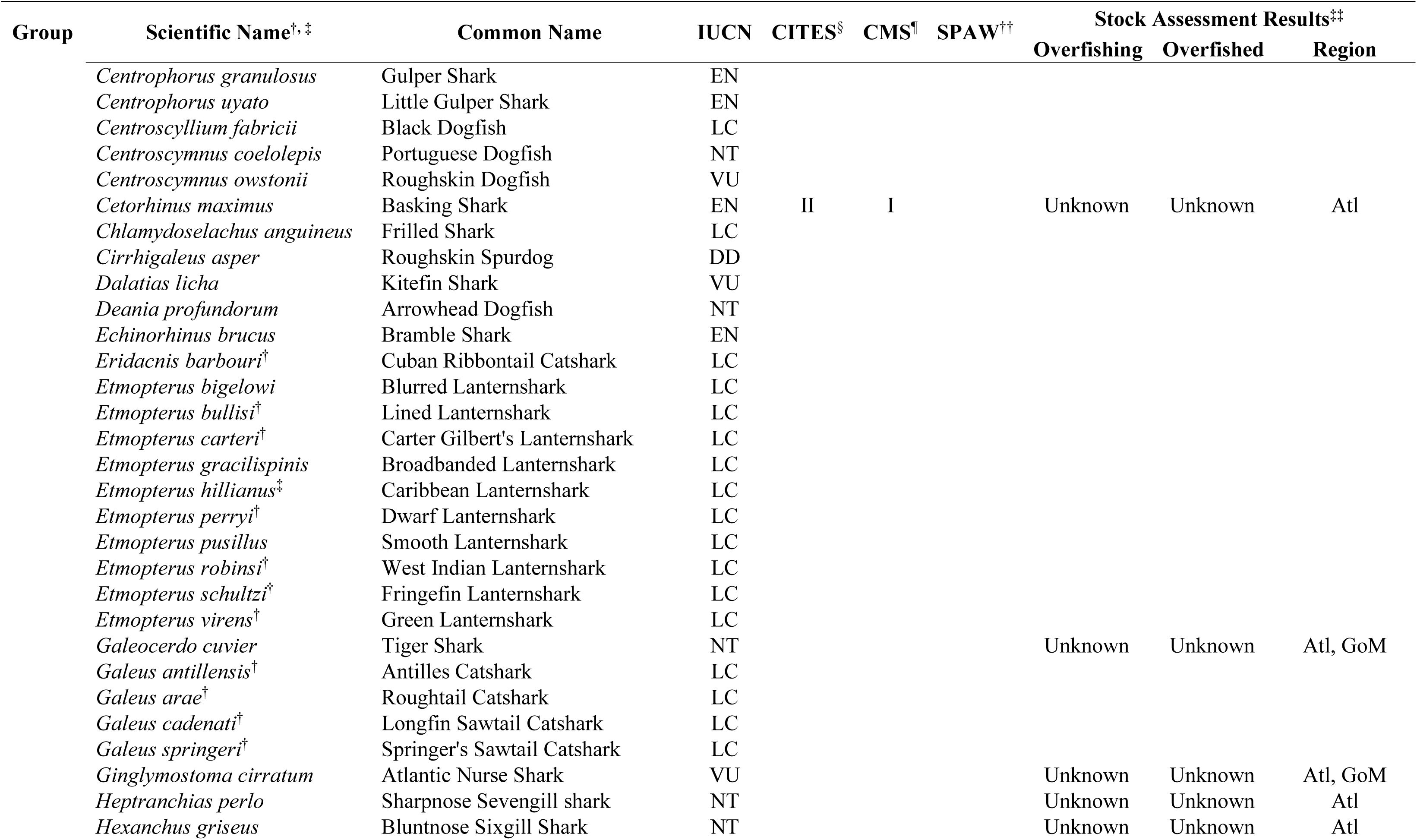
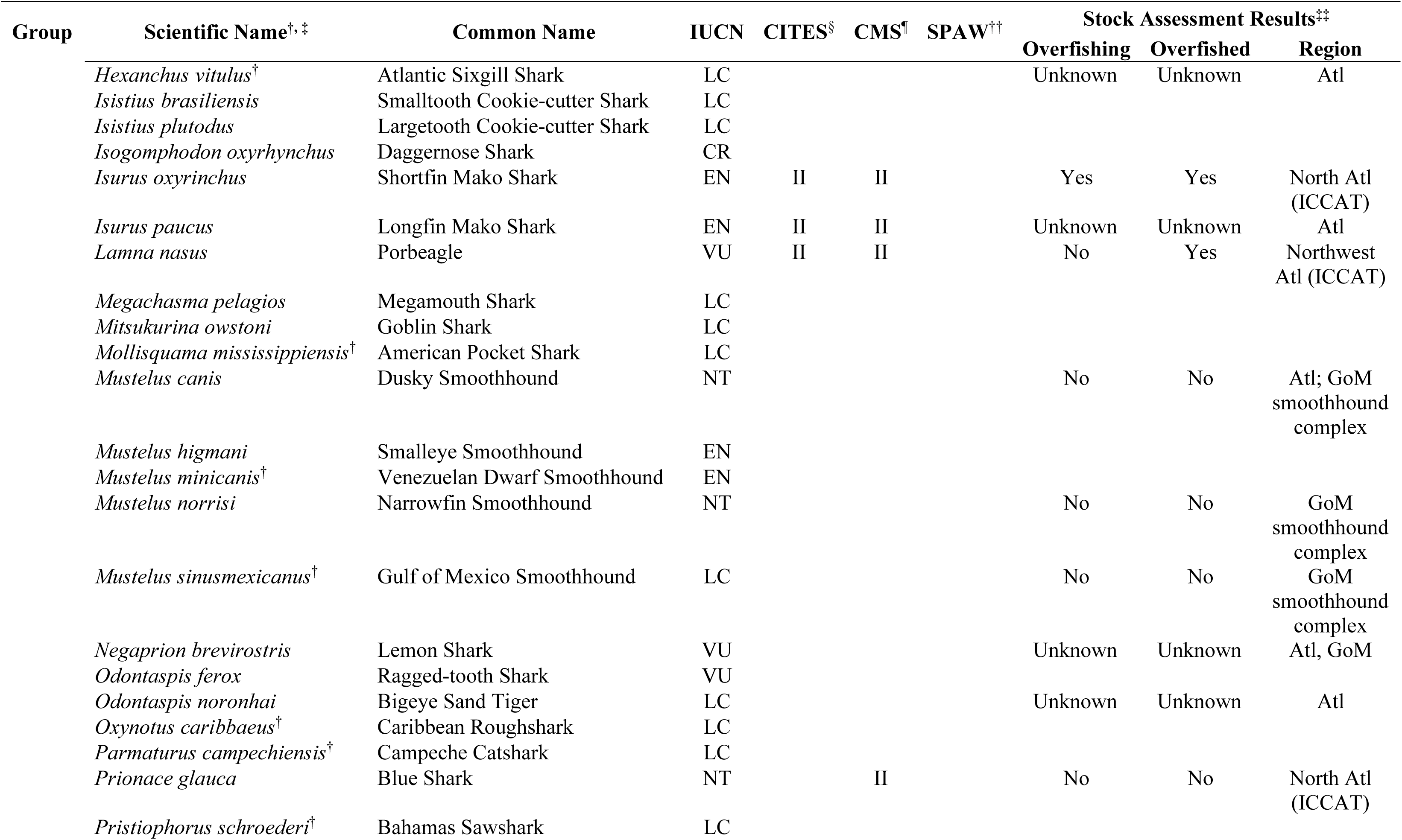
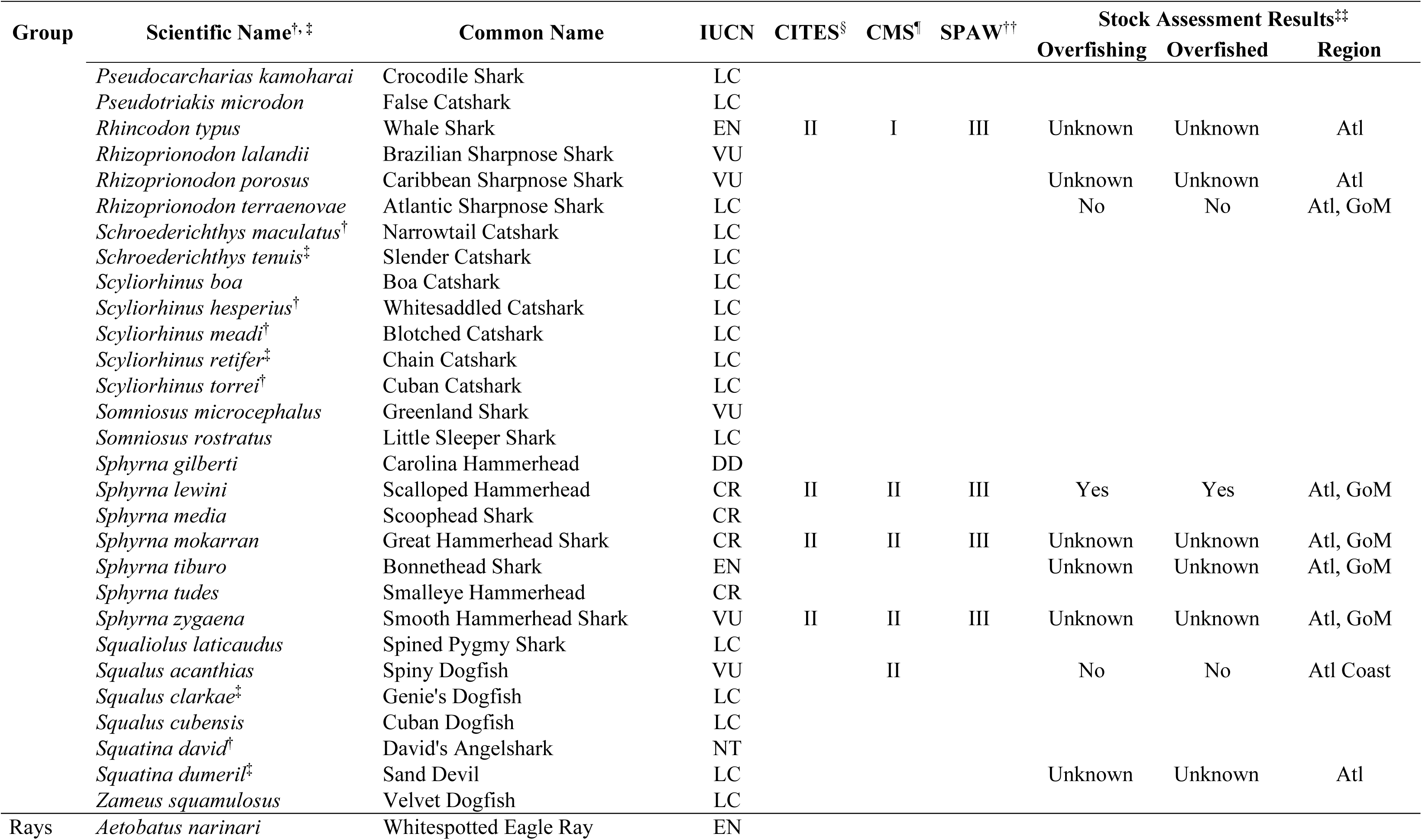
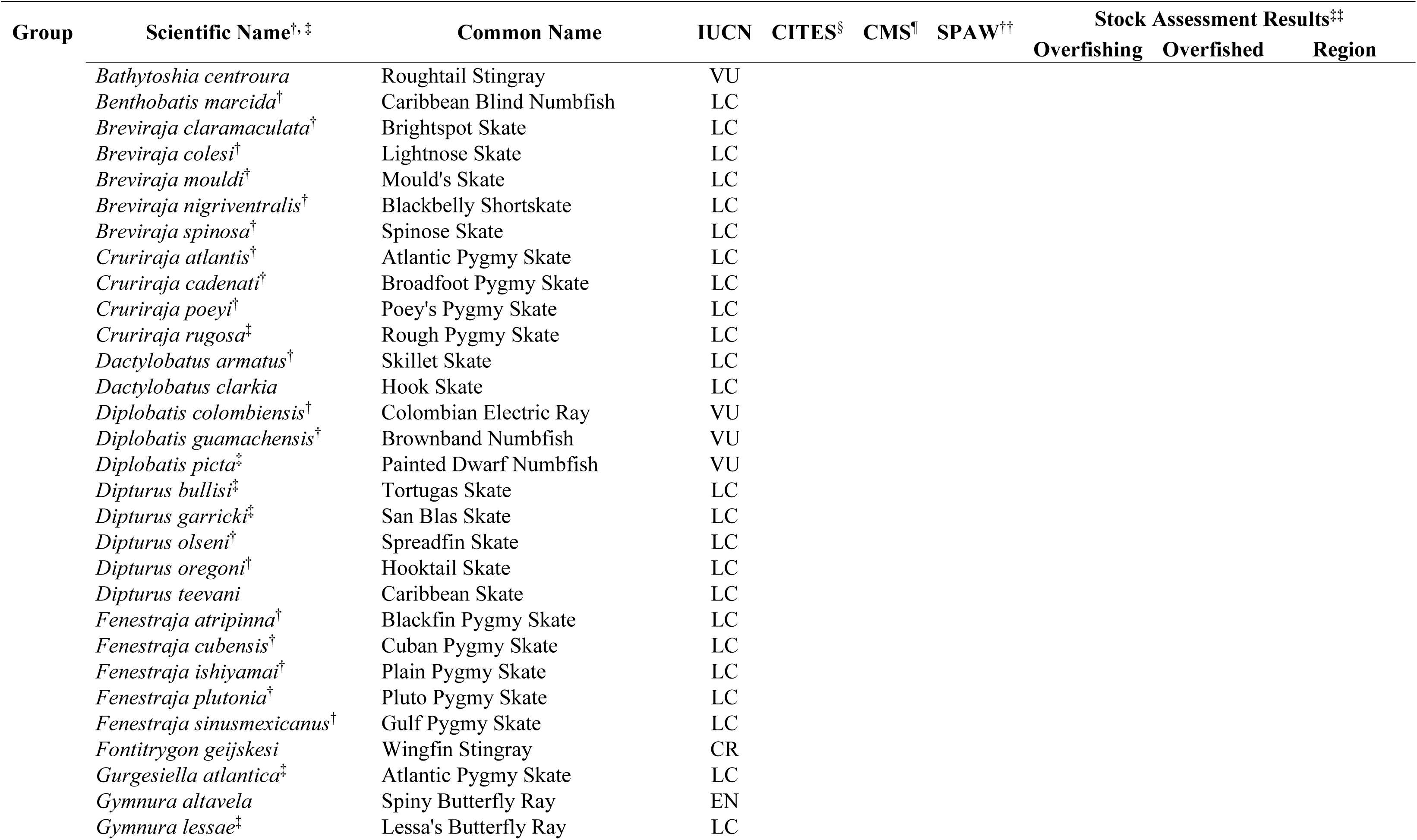
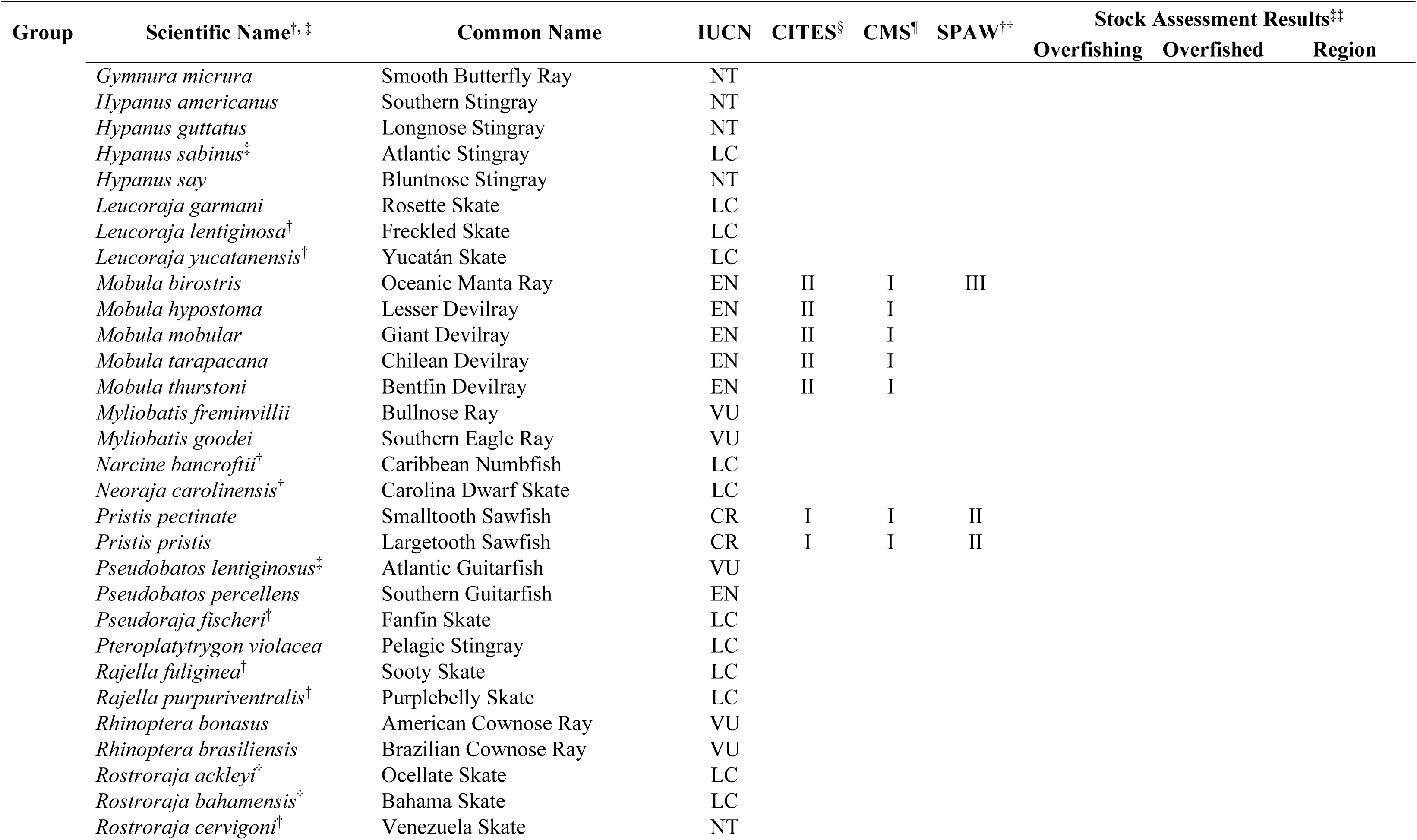
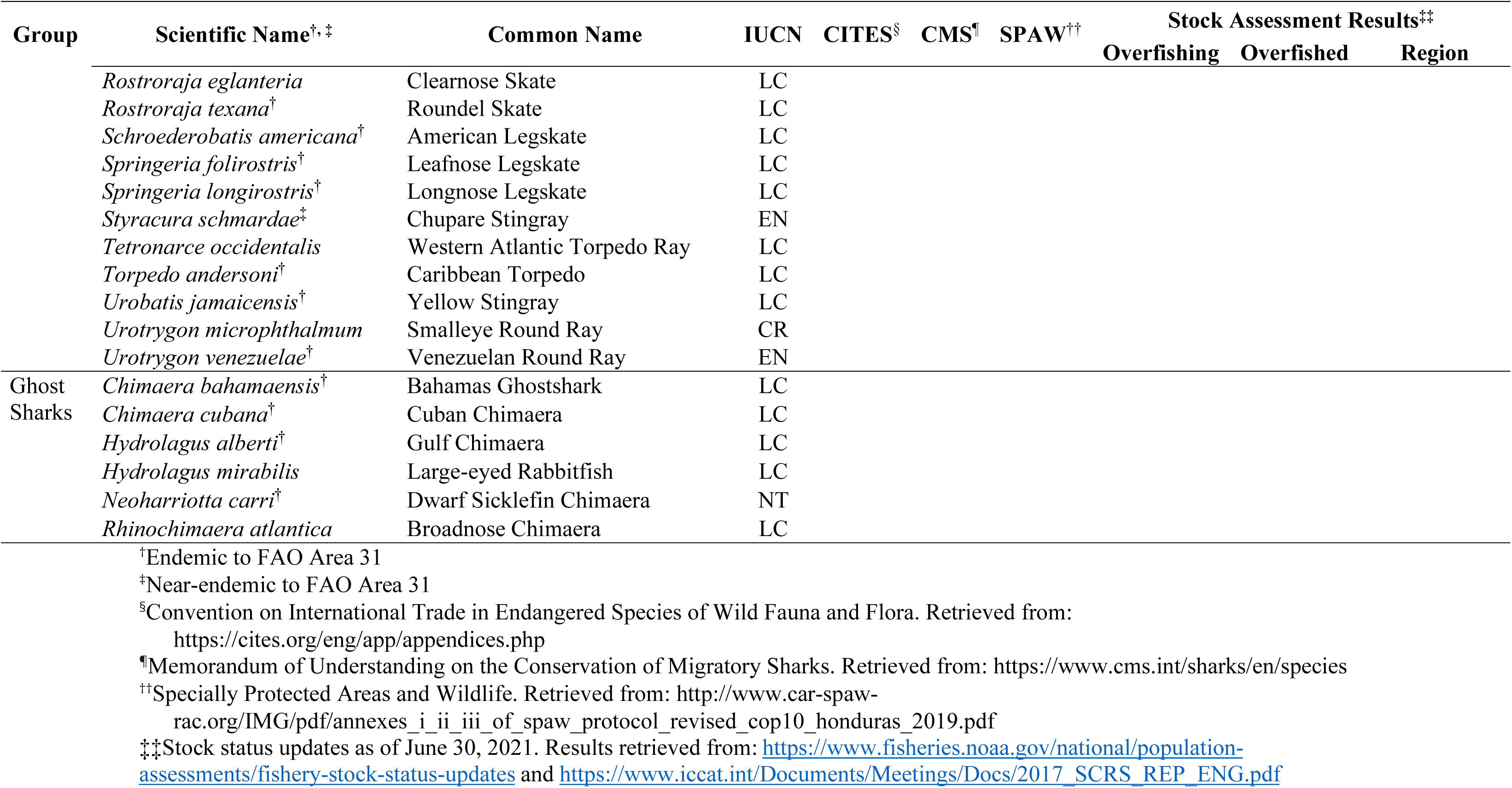
List of chondrichthyans included in this review, including their category of extinction risk according to global assessments by the IUCN, their CITES Appendix, CMS Appendix, and SPAW Annex status, and results of stock assessments according to the United States or ICCAT by region (where ‘Atl’ is Atlantic and ‘GoM’ is Gulf of Mexico). Overfishing refers to fishing mortality being higher than it is at maximum sustainable yield and overfished refers to a stock having a low population size that threatens its ability to reach maximum sustainable yield. Species are organized alphabetically within broad taxonomic groups

**Table S2:**
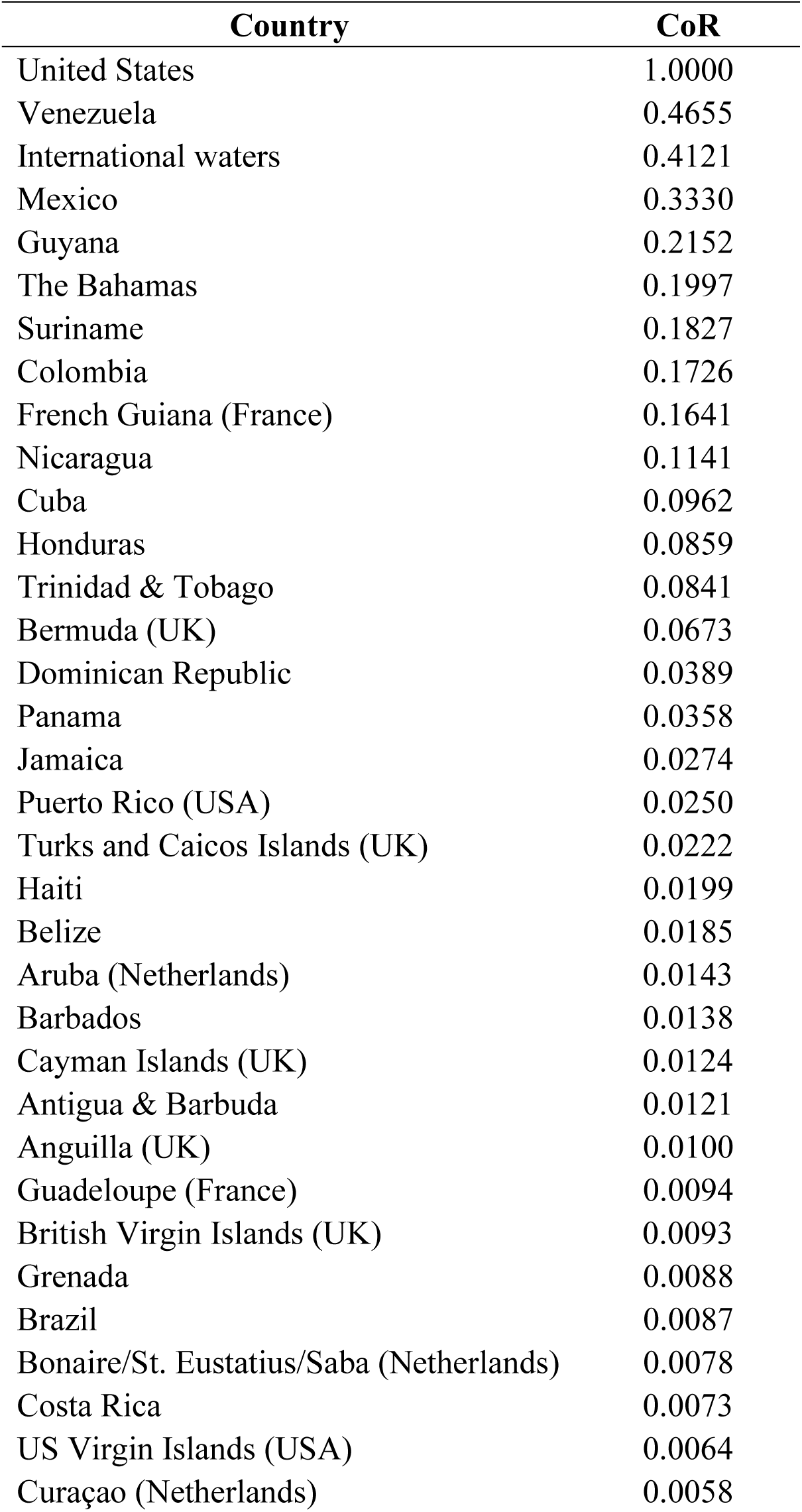
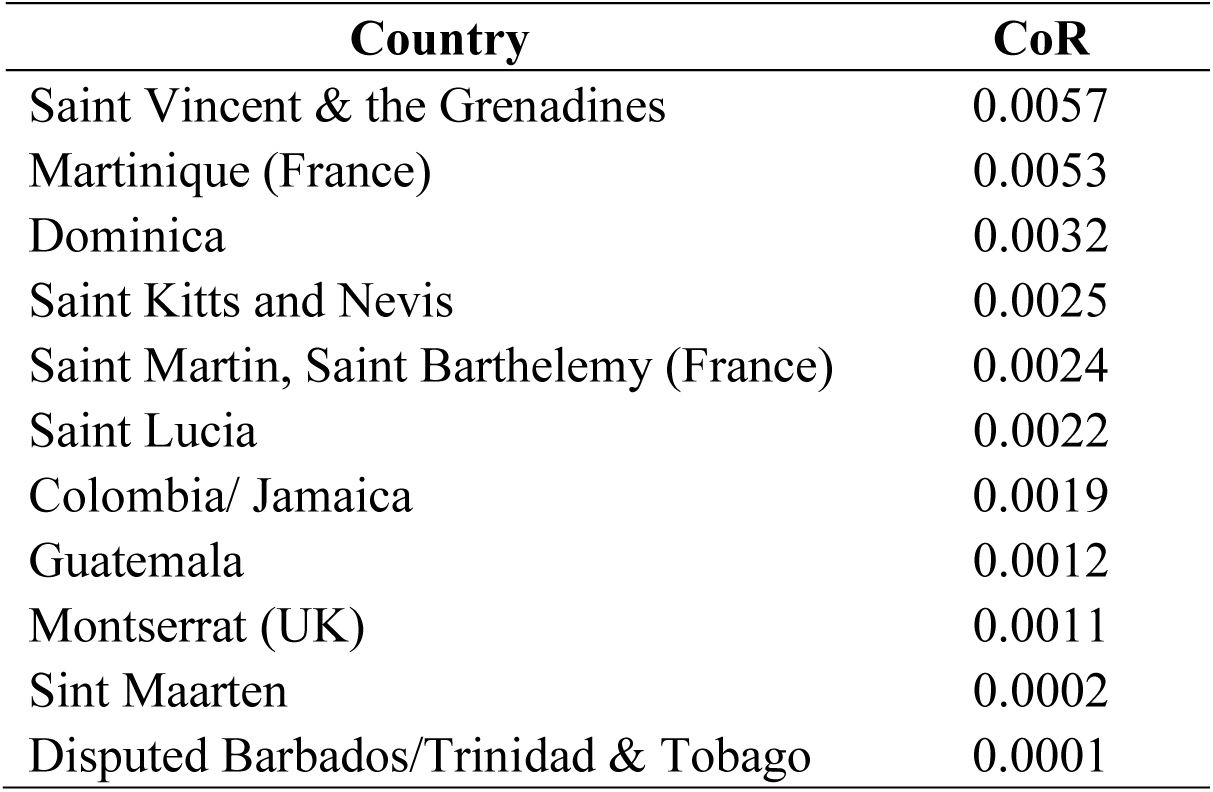
Conservation responsibilities (CoRs) for all chondrichthyans in the Western Central Atlantic Ocean across 44 countries and territories as well as international waters. CoR is a function of extinction risk and proportional species distributions within a given jurisdiction and is normalized from 0 to 1

Table S3: See separate Excel file.

**Table S4:** Chondrichthyan Management Responsibility Scores (‘CMR Score’) for countries in the Western Central Atlantic Ocean. CMR Score is a function of each country’s engagement with thirteen management tools that range from national-level fishing bans to participation in international trade agreements (called Management Engagement), extinction risk and proportional species distributions within a given jurisdiction (called Conservation Responsibility), and total reconstructed catch of sharks and rays from 1950 to 2016. CMR Scores are normalized from 0 to 1, where the highest score (USA) was assigned a 1. The higher the CMR Score, the more unmitigated responsibility to manage sharks and rays. Note that some territories were omitted due to the nature of the underlying data

| Country | Chondrichthyan Management Responsibility Score |
| --- | --- |
| United States | 1.00000 |
| Venezuela | 0.53274 |
| Mexico | 0.34925 |
| Guyana | 0.03942 |
| Suriname | 0.03300 |
| Cuba | 0.03150 |
| Jamaica | 0.01373 |
| French Guiana (France) | 0.01135 |
| Dominican Republic | 0.01055 |
| Trinidad & Tobago | 0.01025 |
| Nicaragua | 0.00933 |
| Colombia | 0.00222 |
| Belize | 0.00178 |
| Barbados | 0.00034 |
| The Bahamas | 0.00025 |
| Honduras | 0.00014 |
| Turks and Caicos Islands (UK) | 0.00010 |
| Martinique (France) | 0.00009 |
| Panama | 0.00006 |
| Bermuda (UK) | 0.00005 |
| Costa Rica | 0.00005 |
| Antigua & Barbuda | 0.00004 |
| Grenada | 0.00003 |
| Aruba | 0.00001 |
| Saint Vincent & the Grenadines | 0.00001 |
| Guadeloupe (France) | 0.00001 |
| Cayman Islands (UK) | 0.00001 |
| Curaçao (Netherlands) | 0.00001 |
| Haiti | 0.00001 |
| Saint Martin, St. Barthelemy, Sint<br>Maarten (France, Netherlands) | 0.00001 |
| Saint Lucia | 0.00000 |
| Bonaire, Sint Eustatius, Saba<br>(Netherlands) | 0.00000 |
| Guatemala | 0.00000 |
| Dominica | 0.00000 |
| British Virgin Islands (UK) | 0.00000 |
| Brazil | 0.00000 |
| Anguilla (UK) | 0.00000 |
| Montserrat (UK) | 0.00000 |
| Puerto Rico (USA) | 0.00000 |
| Saint Kitts and Nevis | 0.00000 |

